# Pathway-based, reaction-specific annotation of disease variants for elucidation of molecular phenotypes

**DOI:** 10.1101/2023.10.18.562964

**Authors:** Marija Orlic-Milacic, Karen Rothfels, Lisa Matthews, Adam Wright, Bijay Jassal, Veronica Shamovsky, Quang Trinh, Marc Gillespie, Cristoffer Sevilla, Krishna Tiwari, Eliot Ragueneau, Chuqiao Gong, Ralf Stephan, Bruce May, Robin Haw, Joel Weiser, Deidre Beavers, Patrick Conley, Henning Hermjakob, Lincoln D. Stein, Peter D’Eustachio, Guanming Wu

## Abstract

Disease variant annotation in the context of biological reactions and pathways can provide a standardized overview of molecular phenotypes of pathogenic mutations that is amenable to computational mining and mathematical modeling. Reactome, an open source, manually curated, peer-reviewed database of human biological pathways, provides annotations for over 4000 disease variants of close to 400 genes in the context of ∼800 disease reactions constituting ∼400 disease pathways. Functional annotation of disease variants proceeds from normal gene functions, through disease variants whose divergence from normal molecular behaviors has been experimentally verified, to extrapolation from molecular phenotypes of characterized variants to variants of unknown significance using criteria of the American College of Medical Genetics and Genomics (ACMG). Reactome’s pathway-based, reaction-specific disease variant dataset and data model provide a platform to infer pathway output impacts of numerous human disease variants and model organism orthologs, complementing computational predictions of variant pathogenicity.

## Introduction

Reactome is an open-source, manually curated, peer-reviewed knowledgebase of human biological pathways and an omics data analysis platform initially focused on normal processes [1–3]. An expanded data model enables annotation of protein variants as participants in disease reactions and pathways [4,5], supporting the intended use of Reactome as a disease mechanism elucidation tool [6].

The ACMG Standards and Guidelines for the interpretation of sequence variants [7] provide the evidence framework to classify variants as benign or pathogenic based on 1) population, 2) computational and predictive, 3) functional, 4) segregation, 5) de novo, 6) allelic, 7) another database-derived, and 8) other data. Criteria for assessing the evidence strength in these eight categories can be supporting (BP1-7) or strong (BS1-4) for benign variants, and supporting (PP1-5), moderate (PM1-6), strong (PS1-4) or very strong (PVS1) for pathogenic variants. Adopting ACMG is the first step towards alignment and exchange of Reactome disease variant annotations with ACMG-compliant variant databases such as ClinGen and ClinVar [8].

Rather than comprehensively cataloging disease variants, Reactome describes the impact of representative protein variants on pathway activity through disease pathway-based, disease reaction-specific functional annotations. Reactome disease variants, whenever possible, cross-reference external open-source resources that provide DNA-level annotations and relevant clinical information: OMIM [9], ClinGen Allele Registry [10], ClinVar [11], and disease-specific databases, such as RettBASE [12] for Mendelian disorders; COSMIC [13], ClinGen Allele Registry [10] or ClinVar [11], and Leiden Open Variation Database (LOVD) [14] for cancer variants.

Reactome adheres to the Human Genome Variation Society (HGVS) protein variant nomenclature [15] when available, with the exception of using 1) one letter amino acid abbreviations; and 2) common protein isoform and cleaved protein fragment names.

Computational variant assessment tools, e.g. SIFT [16], PolyPhen-2 [17], Mutation Assessor [18], and AlphaMissense [19], can give false negative and false positive predictions. For example, pathogenic hotspot mutations PIK3CA H1047L and PIK3CA H1047R [20,21] are assessed as “neutral”, “tolerated”, “benign”, and “ambiguous” by Mutation Assessor, SIFT, PolyPhen-2, and AlphaMissense, respectively. Moreover, the common approximation that all truncating mutations are deleterious is not reliably correct. Reactome’s pathway-based, reaction-specific disease variant annotations can be used to identify and fill in gaps in computational predictions, aiding in the interpretation and modeling of clinically-relevant variants [22].

## Materials and Methods

### Determining the scope of Reactome disease variant curation

Annotation of disease variants in Reactome builds on annotation of wild-type biochemical functions of disease genes. Comprehensive variant lists retrieved from variant databases and PubMed [23] publications (Figure 1), are pruned based on the available functional studies.

**Figure 1.**
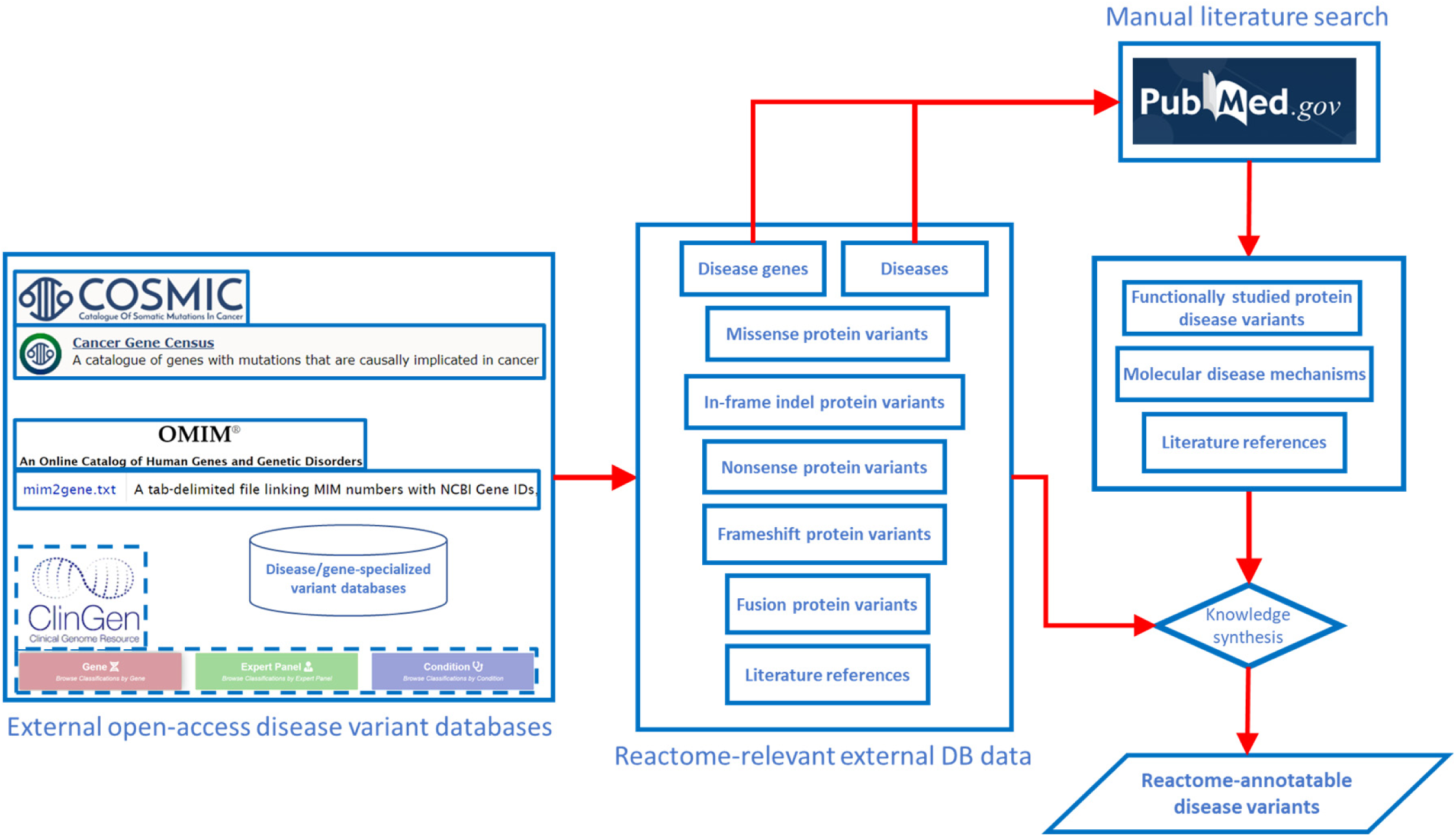
Pipeline for determining the scope of Reactome disease variant curation. ClinGen is shown with dashed borders as it is still to be routinely incorporated in the pipeline.

### Aligning Reactome annotations with ACMG Standards and Guidelines

The ACMG Standards and Guidelines [7] framework for the interpretation of sequence variants based on evidence strength, pathogenicity, and evidence type was evaluated for its ability to describe pathogenicity with Reactome-suitable evidence type and quality level (Table 1).

**Table 1.**
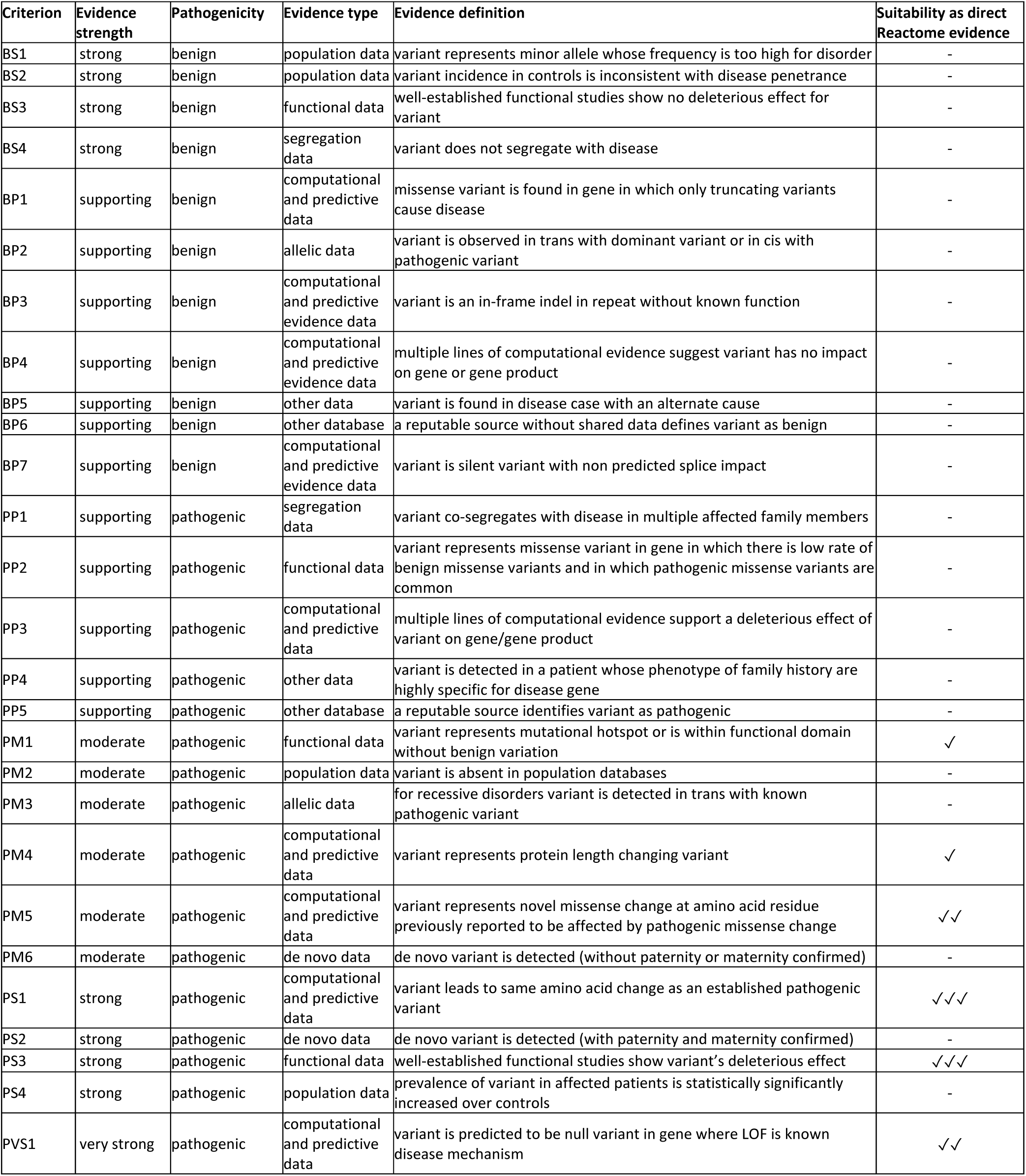
ACMG evidence framework utilized by Reactome. A minus (-) indicates that a criterion cannot be used as direct Reactome evidence. A checkmark (✓) indicates that a criterion can be used as direct evidence, with the number of checkmarks indicating the criterion’s strength from the perspective of Reactome’s data model. This table represents an edited version of previously published Figure 1 [7], reproduced and edited with authors’ and publisher’s permission.

### Reactome data model and curator tool

As previously described [4,5], Reactome annotates disease variants in the context of disease reactions tagged with Disease Ontology (DO) [24] disease terms, and “entityFunctionalStatus” which combines the Sequence Ontology [25] molecular phenotype (e.g. gain_of_function) and sequence variant terms (e.g. missense_variant). Association of the entityFunctionalStatus with disease reactions rather than variants circumvents the problem arising when a mutation differentially affects different functional aspects of a protein [26]. Visualization of disease reactions in the context of normal pathways is achieved by automated overlay onto corresponding wild-type reactions (Figures 2A and 2B) or by direct addition to pathway diagrams (Figure 2C). Functionally analogous variants are grouped into disease variant sets. Detailed description can be found in Supplementary Methods.

**Figure 2.**
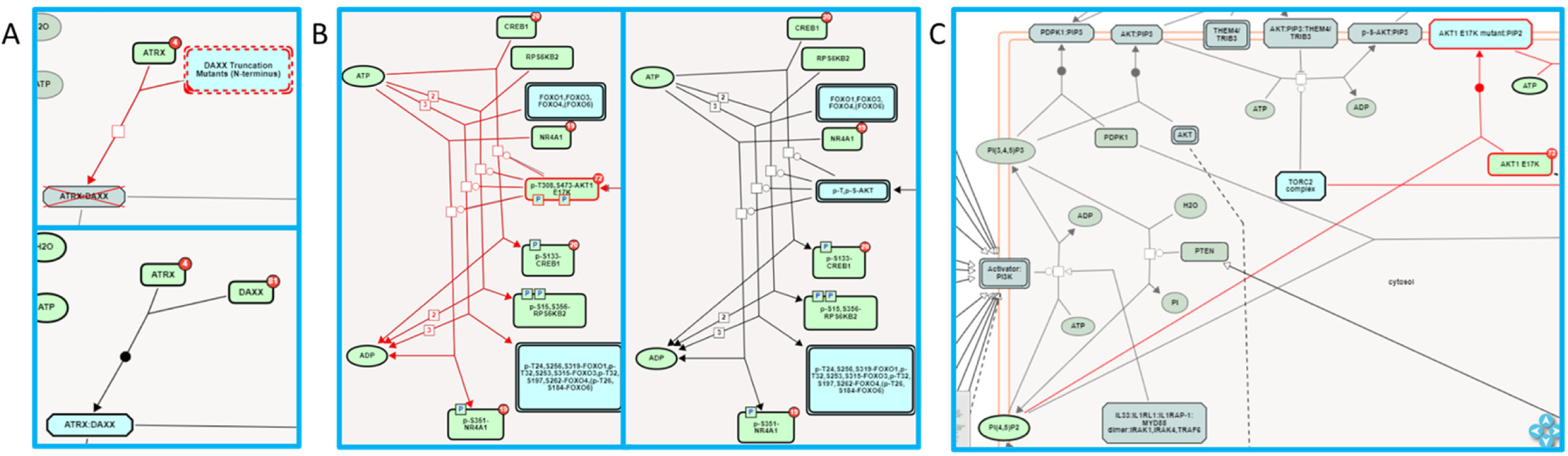
Disease variant-associated disease reactions in Reactome pathway diagrams. **A)** Automated overlay of a LOF reaction “Defective DAXX does not bind ATRX” (upper panel) onto normal reaction “ATRX binds DAXX”(bottom panel) [27]. **B)** Automated overlay of GOF reactions showing phosphorylation of nuclear proteins by oncogenic AKT1 E17K variant (left panel) onto normal reactions in the subpathway “AKT phosphorylates targets in the nucleus” (right panel) [28]. **C)** Manually added GOF reaction “AKT1 E17K mutant binds PIP2”, with no normal reaction counterpart [28].

### Accessing Reactome disease variant data

Reactome disease variant data are stored in the central repository. Only reviewed data are available on Reactome webpages and accessible (https://reactome.org/dev/graph-database#Resources) through a Neo4j Graph Database [29].

Reactome disease variants can be accessed on the Reactome website (www.reactome.org) as previously published [30] and described in the User Guide (https://reactome.org/userguide). Detailed descriptions of annotated disease processes and variants are also available in Reactome’s electronic textbook (https://reactome.org/download/current/TheReactomeBook.pdf.tgz).

### Analysis of disease genes and disease terms in Reactome

Unique HGNC [31] genes with Reactome-annotated disease variants were analyzed for pathway enrichment (Reactome web tool “Analyse gene list function”; Reactome Release 84, March 2023) and overlap with external disease gene databases (BioVenn web tool [32]; COSMIC Cancer Gene Census, COSMIC edition v97, November 2022; ClinGen genes: April 5, 2023 download; OMIM genes: Online Mendelian Inheritance in Man, OMIM®. McKusick-Nathans Institute of Genetic Medicine, Johns Hopkins University (Baltimore, MD), {April 5, 2023}. URL: https://omim.org/). Disease variant-associated DO terms (DO March 2023 release) were analyzed for prevalence and their relationship to the DO tree (constructed using Cytoscape 3.9.1 [33]) was analyzed using PathLinker [34].

## Results

### Disease genes and variants in Reactome

Reactome (version 84 March 2023) includes disease variants for 372 genes (disease genes). Each protein disease variant is annotated as an EntityWithAccessionedSequence (EWAS) instance, characterized by UniProt-derived referenceEntity [35], start and end coordinates, cellular compartment, and a hasModifiedResidue attribute indicating post-translational modifications (PTMs) and mutated residues. One disease variant can have multiple EWASes. Reactome’s disease variant dataset is summarized in Table 2 and provided in Supplementary Table 1.

**Table 2.** Disease variant types in Reactome.

| Variant type | Variant EWASs | Unique variants |
| --- | --- | --- |
| missense | 1955 | 1624 |
| missense in translation initiation codon | 2 | 2 |
| frameshift | 1869 | 1858 |
| nonsense | 1096 | 1087 |
| nonsense leading to usage of an alternative START codon | 1 | 1 |
| fusion | 244 | 122 |
| fusion (fused to promoter of another gene with in-frame deletion) | 1 | 1 |
| in-frame deletion | 91 | 68 |
| in-frame duplication | 36 | 21 |
| in-frame insertion | 33 | 18 |
| in-frame indel | 21 | 12 |
| multiple in-cis mutations | 12 | 9 |
| splicing | 6 | 6 |
| inversion | 2 | 2 |
| extension | 1 | 1 |
| **Total** | **5370** | **4832** |

Overrepresented pathways for Reactome disease genes (Supplementary Table 2, Figure 3), are often related to cancer hallmarks [36] and Mendelian disorders of metabolism, reflecting curation and experimental bias for a small fraction of protein-coding genes involved in particular biological processes [37].

**Figure 3.**
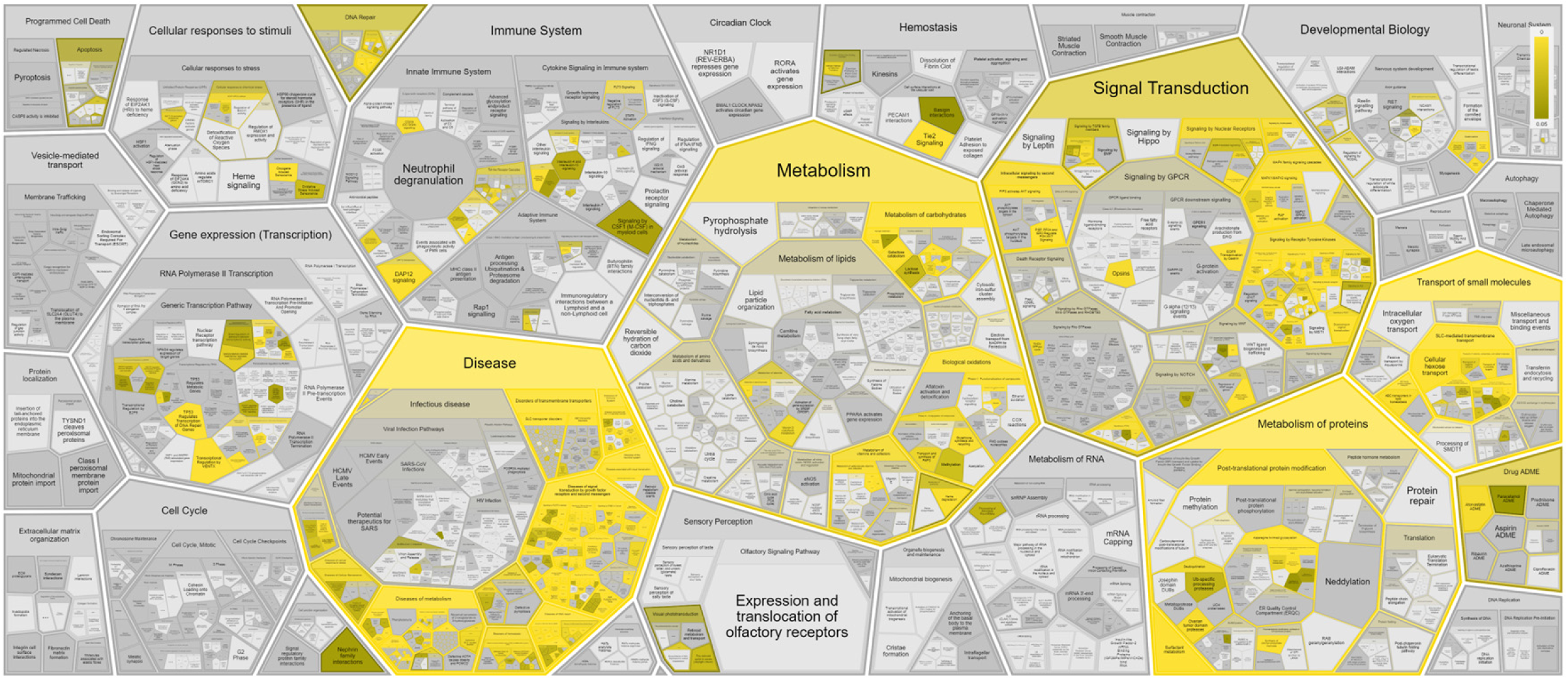
Reactome pathway enrichment analysis results shown in the Reactome Voronoi map for 372 disease variant genes based on Reactome Release 84.

Reactome disease variants cross-reference 421/11254 DO terms (Supplementary Table 3), most frequently from branches shown in Figure 4 and Supplementary Table 4.

**Figure 4.**
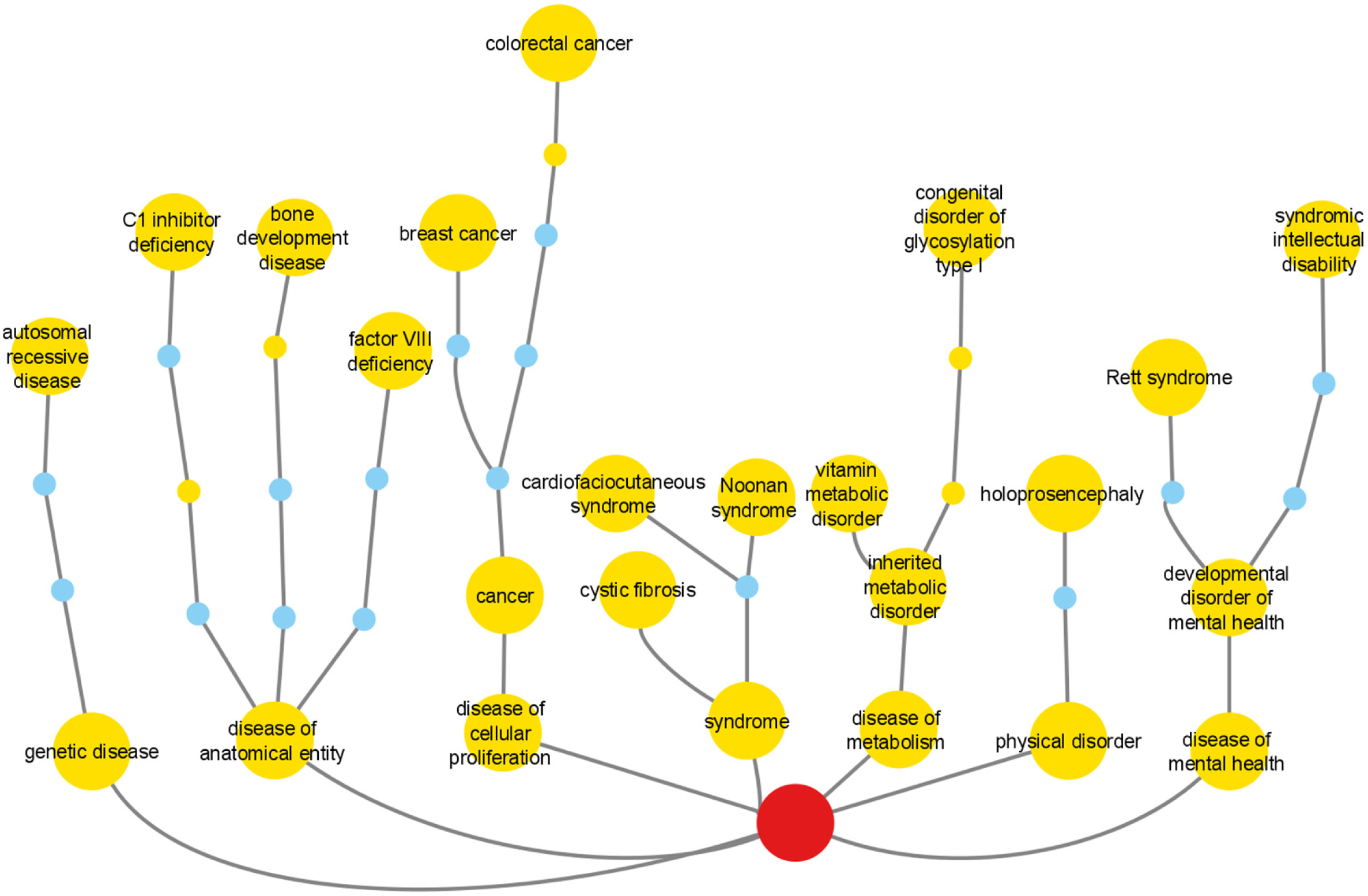
Reactome cross-references to DO branches with most frequently cross-referenced terms for each branch shown. Numbers in bold are the sum values for all cross-references belonging to the branch. Italicized numbers are cross-references to the individual DO term. Yellow nodes are DO terms annotated in the context of Reactome disease variants. Blue nodes are linker nodes, connecting Reactome disease variant-annotated DO terms with the top-level DO term Disease (DOID:4), shown as a red circle.

Reactome variant annotations can lead to novel biological insights by creating explicit connections between disease terms and pathway/process-based molecular phenotypes (Supplementary Table 1), as described in the Discussion.

The overlap of Reactome disease genes with COSMIC v97 Cancer Gene Census List [38], OMIM [9], and ClinGen [39] is shown in Figure 5. Reactome version 84 included 3804 COSMIC, 199 ClinGen, 11 ClinVar, 2 OMIM, and 132 LOVD identifiers. Reactome variant EWASes map to 547 disease variant sets, 801 disease reactions (cross-referencing 506 normal reactions), and 403 diagram-level disease pathways (cross-referencing 73 normal pathways linked to 51 GO Biological Processes) (Supplementary Table 1). The average number of disease reactions a disease gene participates in is 4 (1-54 range, median 1) overall, 8 (1-54 range, median 6) for genes with GOF variants (110/372), 2 (1-16 range, median 1) for genes with LOF variants (262/372). The discrepancy is due to the curation strategy. Reactome disease pathways start with disease reactions diverging from the wild-type and end with disease reactions with all wild-type outputs. With LOF variants, a single disease reaction is frequently the starting and ending reaction in a single disease pathway (e.g. PAH S40L in “Phenylketonuria” [40]). GOF variants frequently engage in a cascade of events (e.g. AKT1 E17K in “PI3K/AKT Signaling in Cancer” [28]).

**Figure 5.**
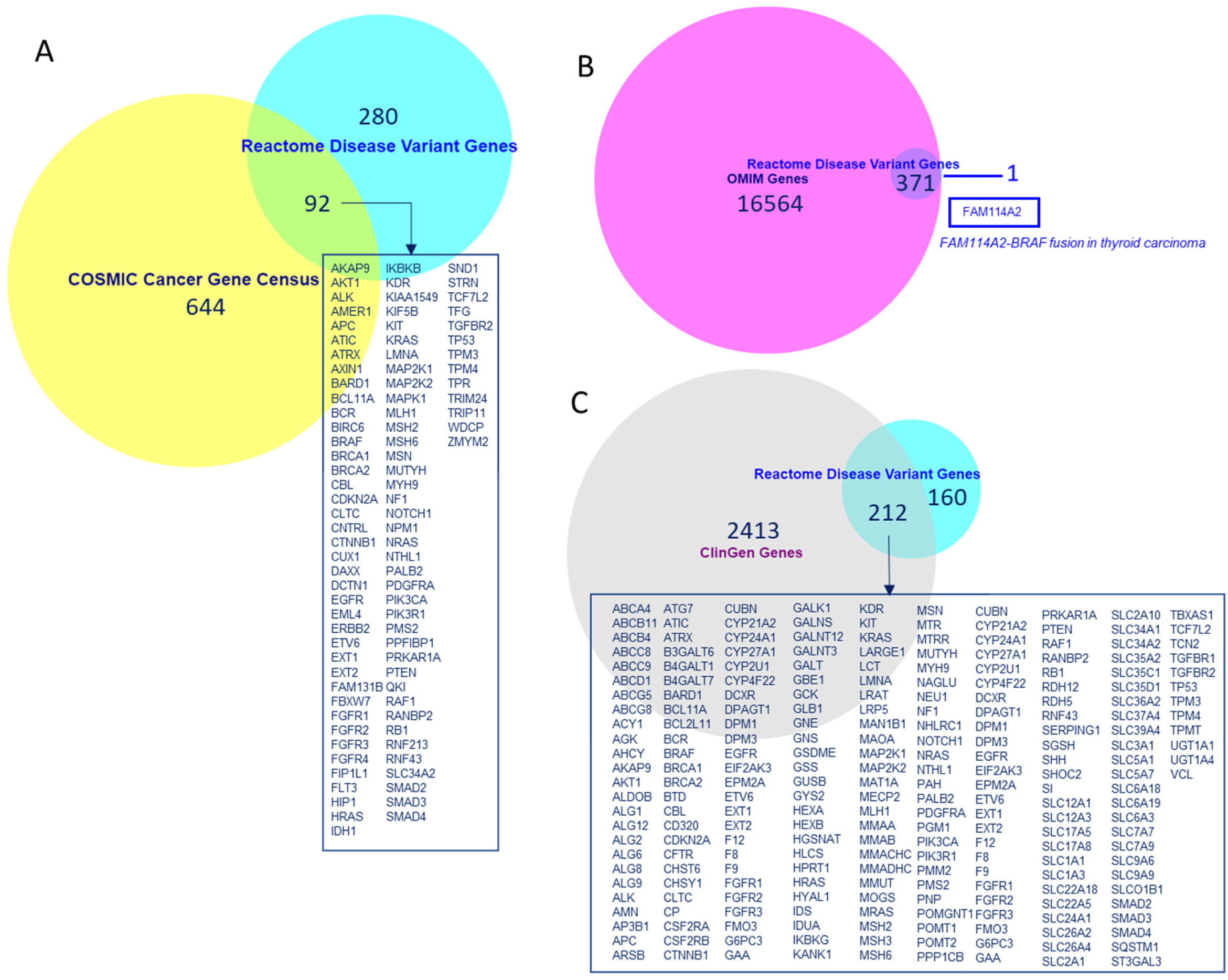
Overlap of Reactome disease variant genes with **A)** COSMIC Cancer Gene Census genes, **B)** OMIM disease genes and **C)** ClinGen disease genes. Overlapping COSMIC/Reactome and ClinGen/Reactome genes are shown. A single disease gene present in Reactome but not in OMIM is shown.

### Reactome-applicable ACMG Standards and Guidelines evidence framework

Two Reactome driving principles are the annotation of disease variants 1) whose molecular functionality has been characterized; and 2) which are clinically relevant/pathogenic.

Benign variants are outside Reactome’s scope. Reactome disease variants correspond to only 6/27 ACMG criteria (PM1, PM4, PM5, PS1, PS3, and PVS1) (Table 1). ACMG criteria BS1-4 and BP1-7 can be used for exclusion of potentially pathogenic variants. ACMG criteria PP1-PP5, PM2, PM3, PM6, PS2, and PS4 are insufficient for reaction-specific annotation but can corroborate pathogenicity of variants conforming to PM1, PM4, PM5, PS1, PS3 or PVS1.

#### PS3, PVS1 and PS1

PS3 variants are the gold standard for Reactome - deleterious effect(s) supported by well-established functional studies - with an additional Reactome requirement for studies to be at the molecular mechanism resolution level. PS3 variants are annotated as functionally analogous members of disease variant sets (e.g. p16INK4 A20P [41–43], Figure 6A) or as direct disease reaction participants (e.g. p14ARF R21Rfs*47 [44,45], Figure 6B).

**Figure 6.**
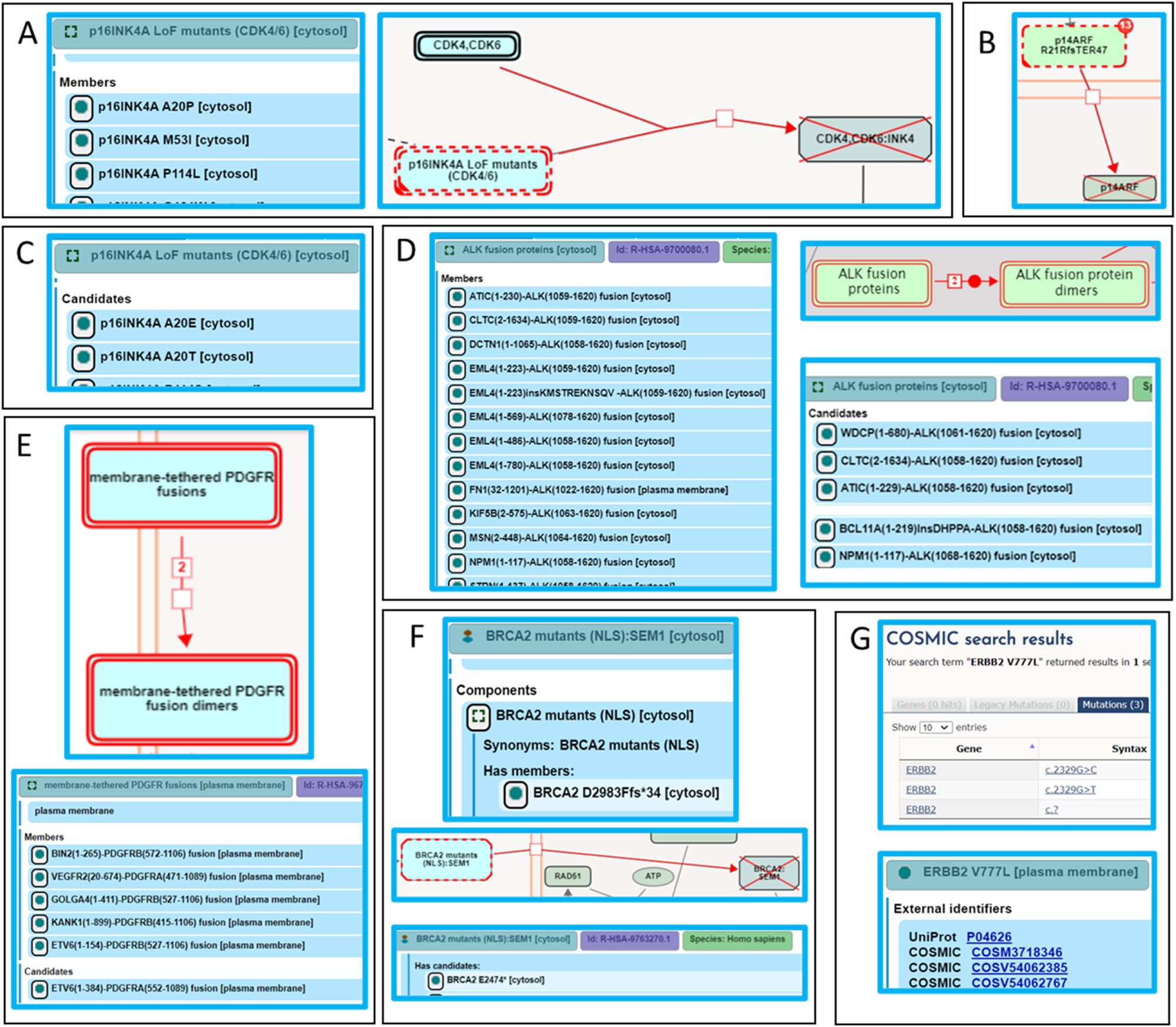
Alignment of Reactome disease variant annotations with ACMG-specified criteria. **A)** PS3 variants are annotated as members of disease variant sets. Cancer variant p16INK4 A20P, unable to bind to and inhibit cyclin-dependent kinases CDK4 and CDK6, is annotated as a member of the “p16INK4A LoF mutants (CDK4/6)” variant set [43]. **B)** PS3 variants are annotated as direct disease reaction participants. Cancer variant p14ARF R21Rfs*47 (p14ARF 60ins16) is unable to translocate to the nucleus [45]. **C)** PM5 variants are annotated as candidates of disease variant sets. Cancer variants p16INK4A A20E and p16INK4A A20T, sharing similarity with the functionally studied p16INK4A A20P, are candidates of the “p16INK4A LoF mutants (CDK4/6)” variant set [43]. **D)** Extrapolation of PS3 and PM5 criteria to fusion variants. Functionally studied (PS3) cancer-associated fusions EML4(1-496)-ALK(1058-1620) and NPM1(1-117)-ALK(1058-1620) undergo ligand-independent dimerization, and are annotated as direct members of the disease set “ALK fusion proteins”. Fusion proteins that have not been functionally characterized, but are expected to behave in a similar manner based on conservation of functional domains in each fusion partner (PM5) are annotated as candidates of the “ALK fusion protein” set [52]. **E)** Further extrapolation of PM5 criterion to fusion variants to include related family members. Fusion variant ETV6(1-154)-PDGFRB(527-1106), known to dimerize independently of ligand stimulation, is a member of the “membrane-tethered PDGFR fusions” variant set, while the analogous, functionally uncharacterized fusion variant ETV6(1-384)-PDGFRA(552-1089), involving PDGFR family member PDGFRA, is a candidate [55]. **F)** Nonsense and frameshift variants not directly functionally studied that conform to both criteria PM1 and PM4 are annotated as candidates of disease variant sets. Cancer variant BRCA2 F2058Lfs*12 is cytosolic and, in accordance with PS3, annotated as a member of the variant set “BRCA2 mutants (NLS)”. Cancer variants BRCA2 E2474*, BRCA2 R2502Lfs*24, and BRCA2 N2553Tfs*95, not functionally studied but similarly lacking the NLS, are annotated as candidates [59]. **G)** Protein disease variants in Reactome cross-reference any PS1 disease DNA variants from relevant reference databases available at the time of annotation. Cancer variant ERBB2 V777L cross-references three applicable COSMIC records: COSV54062385, COSV54062767, and COSM3718346 [46].

Only PVS1 variants that can be annotated at the protein level are included. Unless PS3 criterion is satisfied, PVS1 variants are annotated as disease variant set candidates.

Reactome disease protein variants cross-reference available PS1 DNA variants from reference databases (e.g. ERBB2 V777L [46], Figure 6G).

#### PM5

Reactome extends PM5 variants - novel missense changes at amino acids previously reported to be affected by pathogenic missense changes - to include variants of any type (e.g. fusion, in-frame indel, etc.) that contain mutations analogous to those in PS3 variants as Reactome disease variant set “candidates” (e.g. PM5 variants p16INK4A A20E and p16INK4A A20T of the PS3 variant p16INK4A A20P, Figures 6A and 6C, [43]).

“ALK fusion proteins” set members include EML4(1-496)-ALK(1058-1620), NPM1(1-117)-ALK(1058-1620) [47,48], and others [47,49,50], which undergo ligand-independent dimerization, meeting the PS3 criterion. PM5 candidates of this set include different breakpoints between PS3 fusion protein pairs, e.g. NPM1(1-117)-ALK(1068-1620) [51], or novel fusion partners, e.g. WDCP(1-680)-ALK(1061-1620), containing functional domains analogous to those in PS3 variants (Figure 6D) [52]. In the case of PS3 fusion variant of *PDGFRB*, ETV6(1-154)-PDGFRB(527-1106) [53], a member of the “membrane-tethered PDGFR fusions’’ set, the PM5 criterion was extended to the PDGFR family member PDGFRA to include ETV6(1-384)-PDGFRA(552-1089) [54] as a candidate (Figure 6E) [55]. As no HGVS guidelines exist for the naming of fusion proteins, Reactome proposes nomenclature based on the controlled vocabulary for protein fragments [56], described in Supplementary Methods.

#### PM1 and PM4

Nonsense and frameshift variants not directly functionally studied that conform to both ACMG PM1 (truncation or complete ablation of functional domains key to a particular protein function) and PM4 (protein length changing) criteria are annotated as disease variant set candidates. Of two functional BRCA2 nuclear localization signals NLS1 and NLS2 (positions 3263-3269 and 3381-3385, respectively), only NLS1 is essential [57,58]. BRCA2 F2058Lfs*12 (c.6174delT), common in cancer, is cytosolic [57] and a PS3 member of the “BRCA2 mutants (NLS)” set (Figure 6F). Polymorphisms with intact NLS1, BRCA2 K3326* and BRCA2 E3342*, not cancer-associated (BS2) and properly localized (BS3) [57], are outside Reactome’s scope. Uncharacterized cancer mutants BRCA2 E2474*, BRCA2 R2502Lfs*24, and BRCA2 N2553Tfs*95, with truncations upstream of NLS1, satisfying criteria PM1 and PM4, are annotated as “BRCA2 mutants (NLS)” set candidates [59].

#### Conflicting evidence

When evidence is conflicting, functional data takes precedence in Reactome. If a variant can be categorized as benign by BP4 (computational evidence) and pathogenic by PS3 (functional evidence), as is the case with PIK3CA H1047L and PIK3CA H1047R [20,21], it is annotated as a PS3 variant [28]. If a variant can be categorized as benign by BS3 (functional evidence) and pathogenic by PM5 (computational evidence), it is excluded from Reactome. PS3 variant p16INK4A D74Y, unable to bind and inhibit CDK4 [60] and restrict cellular proliferation [61], has a PM5 variant p16INK4A D74N which retains the ability to bind CDK4 and CDK6 [62] (BS3), although with diminished ability to inhibit cellular proliferation and CDK4/6 catalytic activity [62] (PS3). As the LOF mechanism is uncertain, p16INK4A D74N is waitlisted for future research community contributions [63] and updates [43].

### Retrieval of Reactome disease variant knowledge

Reactome’s disease variant knowledge can be retrieved in three formats: interactive (based on the Reactome’s homepage search function [30]), electronic textbook (https://reactome.org/download/current/TheReactomeBook.pdf.tgz, with heatmap and textbook-style illustration representations [64] shown in Figure 7), and comprehensive tabular (via the Cypher query provided in the Supplementary Methods of the Reactome Graph Database available from the Downloads page, https://reactome.org/download-data, along with Neo4J installation instructions; after Neo4J installation, APOC plugin needs to be installed to run the Cypher query).

**Figure 7.**
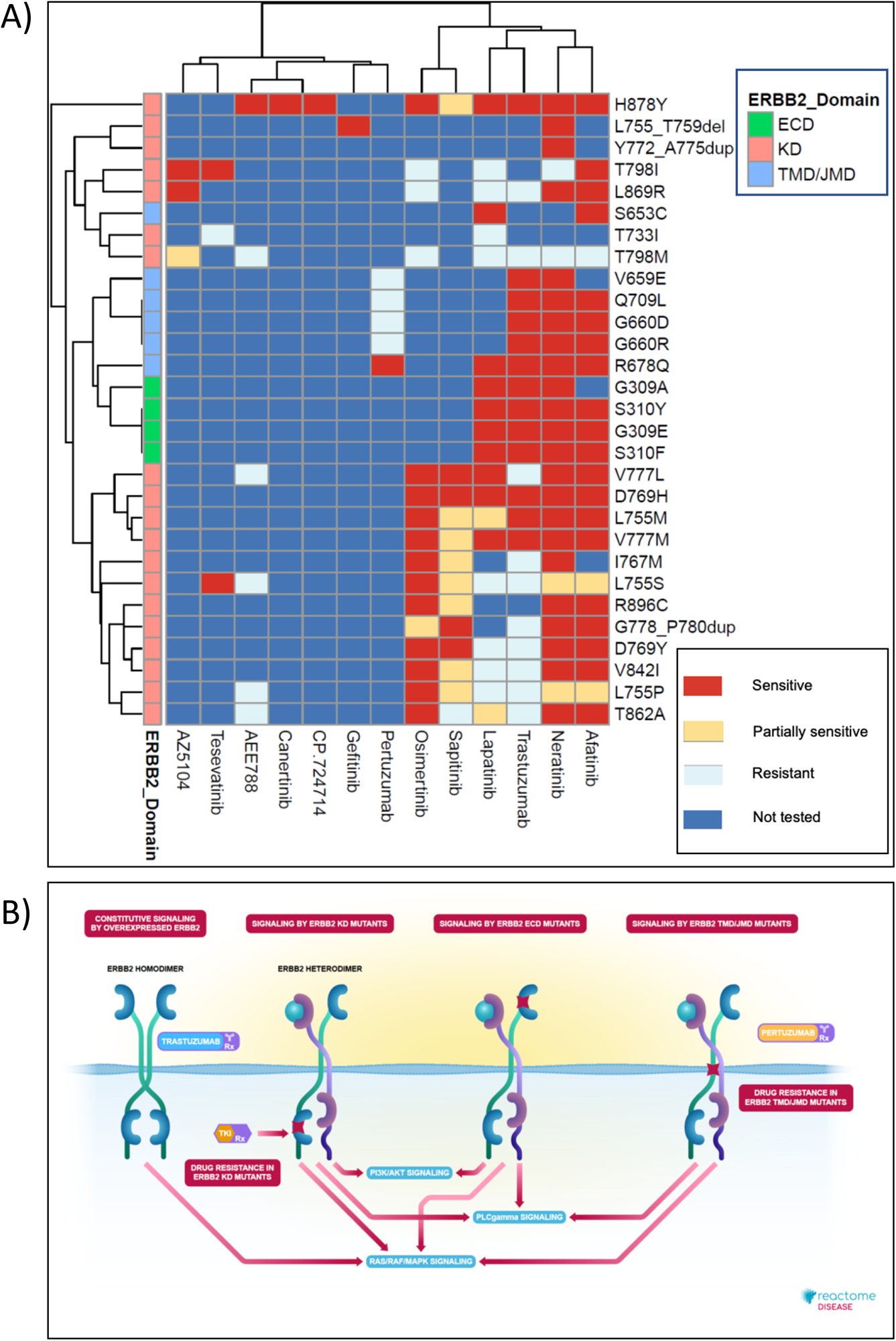
High-level graphical summary of Reactome’s ERBB2 cancer variants content. A) Heatmap representation of Reactome electronic textbook knowledge on the sensitivity of different ERBB2 cancer variants to ERBB2-targeted anti-cancer therapeutics. The heatmap was generated using the R package pheatmap with default settings. B) Interactive textbook style diagram for “Signaling by ERBB2 in Cancer” pathway.

## Discussion

The greatest challenges in predicting pathogenic effects of newly discovered disease-associated variants are the scarcity of mechanistic protein-level evidence, and conflicting predictions by different algorithms. Reactome’s reaction-specific, pathway-based, experimentally supported, peer-reviewed disease variant annotations can be used as a gold-standard dataset for computational pathogenicity assessment based on machine learning or similar approaches, e.g. AlphaMissense [19].

Truncating variants are commonly interpreted as LOF, but mutation position [65] and the surrounding sequence landscape [66] affect the likelihood of nonsense-mediated decay (NMDA). The role of truncated/obliterated protein domains also has to be considered. NOTCH1 PEST domain truncations enhance NOTCH1 oncogenic signaling by interfering with ubiquitin-mediated degradation [67–70].

Many variants of unknown significance (VUS) have been annotated as Reactome disease variant set candidates by extrapolating wild-type protein domain knowledge and experimental findings from PS3 to analogous variants. The Reactome pipeline for bulk automated annotation of pre-selected candidate variants (Supplementary Methods) will enable future automated VUS projection onto Reactome pathways. This requires higher resolution annotations of wild-type protein domains and critical amino acid residues. The large scale, highly accurate protein 3D structures predicted by AlphaFold [71] may facilitate this type of annotation.

Reactome relies on external variant databases and disease/phenotype ontologies for comprehensive clinical information. While continuous communication across databases is necessary for maintaining data integrity, and while open-source databases strive to adhere to FAIR principles [72], lack of standardization hampers FAIRness. Large disease variant repositories do not mandate compliance of disease terms with standardized ontologies, such as DO [24], Monarch Disease Ontology (MonDO) [73] and Human Phenotype Ontology (HPO) [74]. Scientific journals largely do not mandate HGVS nomenclature [15]. Some HGVS guidelines are complex and hard to follow, e.g. nomenclature of initiating methionine missense variants with many synonymous alternatives. For some groups of variants, especially fusion proteins, HGVS guidelines are lacking. Reactome’s fusion protein nomenclature system can improve standardization of fusion protein annotations by HGVS and emerging fusion protein databases FusionGDB [75], CIViC [76], and FPIA [77].

To improve organization of disease variant content and its FAIR compliance, Reactome will provide standardized molecular phenotypes for annotated variants, derived from the entityFunctionalStatus of relevant disease reactions (e.g. LOF or GOF), and the GO Biological Process of the relevant normal pathway counterpart (Supplementary Table 1), thus bridging GO Biological Processes and terms from disease/phenotype ontologies. In addition, Reactome is developing more sophisticated User Guide GraphQL protocols (https://reactome.org/userguide), planning a disease filter for search results, adding the comprehensive disease variant dataset table to its Downloads page, working to systematically apply ClinGen/ClinVar cross-references to all applicable disease variants along with ACMG pathogenicity criteria tags, developing a protocol for contributing its disease variant annotations to open-source disease variant databases, and improving its pipeline for periodic updates of external cross-references.

## Acknowledgements

We would like to acknowledge all scientists who critically reviewed or co-authored our disease variant-related content over the years (in alphabetical order): Dr. Rosemary J. Akhurst, Dr. Sandra Alves, Dr. Mahdi Amiri, Dr. Sanjeevani Arora, Dr. Jane Ashworth, Dr. Richard J. Baer, Dr. Katsiaryna Belaya, Dr. Dorothy C. Bennett, Dr. William Blaner, Dr. Istvan Boldogh, Dr. Ron Bose, Dr. Stefan Broer, Dr. John Christodoulou, Dr. Maria Coutinho, Dr. Richarda M. de Voer, Dr. Frederick Andrew Dick, Dr. Paul W. Doetsch, Dr. Shereen Zarif Ezzat, Dr. Alfonso García-Valverde, Dr. Evripidis Gavathiotis, Dr. Heidi Greulich, Dr. Richard P. Grose, Dr. Lars Hansen, Dr. Nicholas K. Hayward, Dr. Wolf-Dietrich Heyer, Dr. Giorgio Inghirami, Dr. Carman K. M. Ip, Dr. Hiren Joshi, Dr. Rama Krishna Kancha, Dr. Thirumala-Devi Kanneganti, Dr. Julhash U. Kazi, Dr. Anagha Krishna, Dr. Rahul Krishnaraj, Dr. Roland P. Kuiper, Dr. Najoua Lalaoui, Dr. Hang Phuong Le, Dr. Jie Liu, Dr. Yulu Cherry Liu, Dr. Jean-Yves Masson, Dr. Liliana Matos, Dr. Douglas R. McDonald, Dr. Alan K. Meeker, Dr. Dominique Meyer, Dr. Larissa Milano de Souza, Dr. Karobi Moitra, Dr. Gemma Montalban, Dr. Hassan Y. Naim, Dr. Yusaku Nakabeppu, Dr. Toshio Nakaki, Dr. Vaishnavi Nathan, Dr. Bart Pederson, Dr. Daniel Pilco-Janeta, Dr. Steven W. Polyak, Dr. Helmut Pospiech, Dr. Roger R. Reddel, Dr. Barbara Rivera Polo, Dr. Helen Rizos, Dr. Mark G. Rush, Dr. Kazuyasu Sakaguchi, Dr. Sima Salahshor, Dr. Harini Sampath, Dr. Sevtap Savas, Dr. César Serrano, Dr. Feng Shao, Dr. Suneet Shukla, Dr. Dorothe Spillmann, Dr. Robert M. Stephens, Dr. Lauren Thorpe, Dr. David J. Timson, Dr. Dean R. Tolan, Dr. Spiros Vlahopoulos, Dr. D Watkins, Dr. Robert Winqvist, Dr. Jim Woodgett, Dr. Haluk Yuzugullu, Dr. Bin Zhang, Dr. Zhibin Zhang, Dr. Jean J. Zhao, Dr. Jia Zhou.

