## Supplementary Methods for "Pathway-based, reaction-specific annotation of disease variants for elucidation of molecular phenotypes"

**Affiliations**
[1] Adaptive Oncology, Ontario Institute for Cancer Research, Toronto, ON, Canada, M5G 0A3
[2] New York University Grossman School of Medicine, New York, NY 10016, USA
[3] Oregon Health and Science University, Portland, OR 97239, USA
[4] European Molecular Biology Laboratory, European Bioinformatics Institute (EMBL-EBI), Wellcome Genome Campus, Hinxton, Cambridgeshire, CB10 1SD, UK
[5] College of Pharmacy and Health Sciences, St. John's University, Queens, NY 11439, USA
[6] Open Targets, Cambridge, UK
[7] Department of Molecular Genetics, University of Toronto, Toronto, ON, Canada, M5S 1A8
[*] Currently: Data Curation and Modelling Data Office, Data Science and AI, AstraZeneca UK, Academy House, Cambridge, Cambridgeshire, CB2 8PA, UK

### Supplementary Methods

#### Guidelines for manual and automated functional annotation of disease variants

Depending on the scope and divergence of pathogenic variants of a particular disease gene, Reactome applies procedures for either manual or automated functional annotation of disease variants. Manual annotation of disease variants is appropriate when the number of representative variants is small and when representative variant types are dissimilar, as is the case, for example, with ERBB2, NOTCH1 and CFTR variants (Supplementary Table 1). Automated annotation is appropriate when the number of representative variants of the same variant type is large, as is the case, for example, with RB1 and TP53 variants.

Manual annotation of disease variants in Reactome proceeds through the following steps (Supplementary Methods Figure 1A):

1) Disease gene prioritization
2) Mapping molecular disease mechanisms to wild-type (normal) Reactome pathways
3) Defining disease pathways and, if possible, how they will be linked to normal pathways
4) Defining disease reactions representative of disease molecular mechanism and, if possible, how they will be linked to normal reactions
5) Defining disease variant participants of disease reactions and their classification as direct participants or members/candidates of disease variant sets
6) Defining entity functional status of disease reactions
7) Creation of interactive disease pathway diagrams

For a detailed protocol, please refer to “General approaches to disease entity annotation and naming in Reactome” below, an excerpt from Reactome’s Curator Guide.

#### Guideline for automated annotation of disease variants

Automated annotation of disease variants in Reactome proceeds through the following steps (Supplementary Methods Figure 1B):

1) Starting from manually annotated disease variants grouped into functional sets, define external database-derived and/or literature-derived variants with matching disease association that conform to Reactome-expanded ACMG criterion PM5 or to combined ACMG criteria PM1 and PM4.
2) Categorize variants of interest by mutation type (missense, nonsense, etc.).
3) Run the script that creates individual disease variant records for automatic upload to the Reactome Curator Tool.
4) Import automatically annotated disease variants into existing disease variant functional sets.
5) Expanded disease variant functional sets are readily displayed in interactive pathway diagrams.

#### General approaches to disease entity annotation and naming in Reactome

**Naming:** Reactome variant protein names are formulated in accordance with the HGVS nomenclature [1], with the exception of using single letter abbreviations for amino acid residues. When a descriptor is added (usually to set names), ‘variant’ is to be used rather than ‘mutant’ as in “PI3K variants”. Specific rules for the naming of different variants are described more fully in the relevant sections below.

**Overview:** When creating a database record for a disease variant, it is often most efficient to make a copy of the wild-type protein record, change the relevant fields, and add the disease-specific attributes. Like disease events, disease entities must be labeled with a disease tag, currently derived from the Disease Ontology (DO). Where possible, disease entities will have a cross-reference to an external database such as COSMIC, OMIM, or ClinGen. Disease entities should also have their literature reference field filled in citing the work where the variant was identified or characterized unless they are annotated as disease variant candidates (Supplementary Methods Figure 2A).

#### Data model classes for ‘hasModifiedResidue’

The changes in sequence that lead to the formation of a variant protein are captured by the field “hasModifiedResidue” using the modified residue class “GeneticallyModifiedResidue” to distinguish genetically encoded variants from post-translational modifications (TranslationalModifications class) and RNA editing (TranscriptionalModification class). Subclasses of the GeneticallyModifiedResidue class are shown in Supplementary Methods Figure 2B. **Note that in both its variant naming convention and in the annotation of**

**‘GeneticallyModifiedResidue’, Reactome focuses on amino acid (protein level) changes and does not capture changes at the nucleic acid level.**

#### Annotation strategies for common classes of protein variants.

##### Missense variants

These variants are the consequence of the replacement of a single amino acid codon with a non-synonymous amino acid codon without change in the reading frame.

**Naming format:** “prefix” + “wild-type amino acid” + “position” + “new amino acid”,

where “prefix” is the HGNC gene symbol and the amino acids are represented by single letter codes.

**Example:** MTR P1173L - substitution of proline 1173 with leucine.

**How to annotate:** Missense mutations are captured with the “ReplacedResidue” subclass of GeneticallyModifiedResidue. PSI-MOD terms [2] are used to construct the replaced residue, identifying both the normal residue with an [amino acid removal] instance and its replacement with another [amino acid residue] instance. Here, L-proline is replaced by L-leucine at coordinate 1173 (Supplementary Methods Figure 2C).

##### Nonsense variants

Nonsense variants are the consequence of mutations that replace a single amino acid codon with a nonsense codon.

**Naming format:** “prefix”+ “amino acid” + “position” + “*”,

where “prefix” is the HGNC gene symbol, the amino acid is represented by a single letter code and the stop codon is represented by an asterisk (*).

**Example:** BRCA2 R2318* - arginine 2318 replaced with stop codon

**How to annotate:** Nonsense mutations are captured using the “NonsenseMutation” subclass of “ReplacedResidue”. A single PSI-MOD term is entered in the record indicating the removal of the wild-type amino acid (Supplementary Methods Figure 3A). On the EWAS record, the endCoordinate is changed to be the last amino acid encoded by the variant (one less than the position of the stop codon).

##### In-frame deletion variants

These variants are the consequence of the removal of a stretch of wild-type amino acid codons between the initiator methionine codon and the stop codon without changing the reading frame.

**Naming format:** "prefix" + "first deleted amino acid" + "first deleted amino acid position" + "_" + "last deleted amino acid" + "last deleted amino acid position" + "del"

Where "prefix" is the HGNC gene name and the deleted amino acids are represented by single letter codes separated by underscore.

Example: HK1 H577_C672del: a deletion of residues H577 to C672, inclusive

**How to annotate:** Deletions are captured using the FragmentDeletionModification class specifying the start and stop coordinate of the deleted stretch of amino acids. In the corresponding EWAS record, the end coordinate of the disease variant protein is not adjusted relative to the wild-type protein (Supplementary Methods Figure 3B).

For deletion of a single amino acid, the start and end position of the FragmentDeletionModification are the same, and the EWAS is named without a range of amino acids, for example, CD320 E88del (Supplementary Methods Figure 3C).

**In-frame insertion variants**

These variants are the consequence of adding a stretch of the amino acid codons between the initiator methionine codon and the stop codon without changing the reading frame of the protein. The inserted amino acids are NOT a copy of the peptide sequence immediately upstream of the insertion site. If this is the case, these variants are annotated as duplication variants (see below).

**Naming format:** "prefix" + "N-terminal flanking amino acid" + "position of N-terminal flanking amino acid" + "_" + "C-terminal flanking amino acid" + "position of C-terminal flanking amino acid" + "ins" + "inserted_sequence",

where "prefix" is the HGNC gene symbol, the N- and C-terminal flanking amino acids are represented by single letter codes and separated by an underscore, and the inserted sequence is listed as single letter amino acid codes.

**Example:** EGFR D770_N771insNPG - in this variant the three amino acids NPG are inserted between D770 and N771. The wild-type protein sequence in this region is 770-DNPHV-774 and the variant sequence is 770-D**NPG**NPHV-. NPG (shown in bold) is inserted at position 771.

**How to annotate:** Insertions are captured using FragmentReplacementModification class specifying the start and stop coordinates which overlap with positions of the N-terminal and C-terminal flanking amino acids, respectively, and specifying the alteredAminoAcidFragment, so that that the first letter in the string is the one letter code for the N-terminal flanking amino acid, the last letter in the string is the one letter code for the C-terminal amino acid, and the string in between corresponds to one letter codes of inserted amino acids. In the EWAS record, the end coordinate of the in-frame insertion variant is not changed compared to the wild-type protein annotation to accommodate the insertion (Supplementary Methods Figure 2D).

**In-frame duplication variants**

These variants are the consequence of in-frame insertion of a stretch of amino acid codons between the initiator methionine codon and the stop codon that ARE synonymous to the coding sequence immediately upstream of the insertion site, leading to a duplication of an amino acid string.

**Naming format:** “prefix” + “first amino acid in the duplicated string”+ “position of the first amino acid in the duplicated string” + “_” + “last amino acid in the duplicated string” + “position of the last amino acid in the duplicated string” + “dup”,

where “prefix” is the HGNC gene symbol, and the amino acids are represented by single letter codes.

**Example:** KIT A502_Y503dup: a duplication of residues A502 and Y503. In this example, wild-type KIT sequence is 502-AYFN-505 and the variant sequence in 502-AY**AY**FN-, with duplicated residues shown in bold and located at positions 504 and 505 relative to the wild-type sequence.

**How to annotate:** Duplications are captured using the FragmentInsertionModification class specifying the start and end coordinates of the duplicated stretch of amino acids in the reference sequence (startPositionInReferenceSequence and endPositionInReferenceSequence, respectively) and with the coordinate of the insertion, which equals endPositioninReferenceSequence+1. As is the case with in-frame deletions and in-frame insertions, the end coordinate of the duplication variant EWAS is NOT changed relative to the wild-type protein (Supplementary Methods Figure 4A). Notice that the referenceSequence of the FragmentInsertionModification is the same as the referenceEntity of the disease variant EWAS. This is in contrast to the annotation of fusion proteins using FragmentInsertionModification, where its referenceSequence differs from the disease variant EWAS reference entity, as described below.

**Fusion proteins**

Fusion proteins are the consequence of in-frame fusion of coding sequences from two different genes, usually through chromosomal translocation. By convention, Reactome annotates these fusions as the insertion of a stretch of the C-terminal partner, using the FragmentInsertionModification class, into the N-terminal partner.

**Naming format:** “prefix (N-terminal protein)” + “(included amino acids from N-terminal protein) + “-” + “prefix (C-terminal protein)” + “(included amino acids from C-terminal protein)” + “fusion”.

Where “prefix” is the HGNC gene symbol and (included amino acids) represents the start and end coordinates of each fusion partner relative to their wild-type reference sequences, in accordance with Reactome’s controlled vocabulary for naming of protein fragments [3].

**Example:** ZMYM2(1-913)-FGFR1(429-822) fusion - this is the insertion of FGFR1 residues 429-822 (relative to full-length wild-type FGFR1) after residue 913 of ZMYM2.

**How to annotate:** Fusions are captured as “FragmentInsertionModifications” class, specifying the start and end coordinates of the C-terminal partner relative to the wild-type reference sequence. In the EWAS record, the end coordinate of the N-terminal partner is changed to the last amino acid of the N-terminal partner sequence that is contained in the fusion (Supplementary Methods Figure 4B). Post-translational modifications to either partner in the fusion protein are numbered according to each respective wild-type reference protein sequence and do not reflect the amino acid positions in the fusion. For instance, if in the context of the fusion protein, the FGFR1 partner is phosphorylated at the reference FGFR1 position Y766, the fusion protein would be ZMYM2-p-Y766-FGFR1, even though in the linear sequence the phosphorylation occurs at residue 1251 of the fusion (913 + 766 - 428).

**If a fusion protein includes an in-frame insertion of amino acids encoded by an intronic or UTR sequence** that are not present in the reference protein sequences of either of the two fusion partners, the insertion is defined in the name of the fusion protein in the format:

“prefix (N-terminal protein)”+ “(included amino acids from N-terminal protein)” + “ins” + “inserted sequence” + “-” + “prefix (C-terminal protein)” + “(included amino acids from C-terminal protein)”+ fusion,

where the inserted sequence is listed as single letter amino acid codes.

**Example:** FIP1L1(1-391)insAQCP-PDGFRA(577-1089) fusion - this is an insertion of the amino acid string AQCP between the amino acid 391 of FIP1L1 and amino acid 577 of PDGFRA.

**How to annotate:** An in-frame insertion of an amino acid string encoded by an intronic or UTR sequence that is not present in the reference protein sequences of either of the two fusion partners is annotated using the replacedFragmentModification, specifying that the last amino acid of the N-terminal fusion partner (proline at position 391 of FIP1L1 in this example) is replaced by a string of amino acids, where the first amino acid in the string is identical to the last amino acid in the N-terminal fragment, followed by the actual inserted string - P**AQCP** in this example, with the actual inserted string shown in bold (Supplementary Methods Figure 4C).

##### In-frame indels

In-frame Indels are variants that are the consequence of simultaneous insertion and removal of a stretch of amino acids from the wild-type sequence without change in the reading frame.

**Naming format:** “prefix” + “first deleted amino acid” + “position of first deleted amino acid” + “_” + “last deleted amino acid” + “position of last deleted amino acid” + “delins” + “inserted sequence”,

where “prefix” is the HGNC gene symbol, and amino acids, including the inserted sequence, are specified using their single letter codes. If a single amino acid is deleted then only that amino acid and its coordinates are specified:

“prefix” + “deleted amino acid” + “position of deleted amino acid” + “delins” + “inserted sequence”

**Example:** KIT T417_D419delinsY - this is removal of residues T417 to D419 from KIT and their replacement with tyrosine residue.

**How to annotate:** In-frame indels are captured with the FragmentReplacedModification class, specifying the start and end coordinates of the deleted amino acid stretch, and specifying the inserted amino acid stretch that replaces them as an alteredAminoAcidFragment, using single letter amino acid codes. The end coordinate of the variant protein's EWAS record is not changed relative to the wild-type protein (Supplementary Methods Figure 4D).

**Frameshift variants**

Frameshift variants are the consequence of mutations that lead to the insertion or deletion of nucleotides where the total number of inserted or deleted nucleotides is not a multiple of three, leading to a change in the reading frame. The published literature generally provides an exact description of the mutation at the DNA level, from which the curator may need to infer the altered C-terminal sequence of the mutated protein to create a "FragmentReplacedModifcation"

**Naming format:** "prefix" + "first amino acid altered due to frameshift" + "position of the first altered amino acid" + "fs" + "*" + "position of the stop codon relative to the first altered amino acid",

where "prefix" is the HGNC gene symbol, single letter amino acid abbreviations are used, "fs" indicates frameshift, and asterisk indicates the stop codon.

**Example:** EXT1 L490Rfs*9 - in this protein, a frameshift mutation causes substitution of L490 with R, and is followed by a novel stretch of 7 amino acids (8 altogether, counting from the first substituted amino acid) before a stop codon is reached at the 9th position.

**How to annotate:** Frameshifts are captured using the FragmentReplacedModification class, specifying the start and end coordinates of the altered sequence, where the startPositionInReferenceSequence matches the position of the first amino acid substituted due to frameshift (490 in this example) and the endPositionInReferenceSequence is calculated by the formula:

endPositionInReferenceSequence = startPositionInReferenceSequence + frameshift length - 2

The frameshift length is specified in the name of the frameshift variant (the number after the asterisk and equals the length of the substituted amino acid string + 1 (as it takes into account the stop codon).

In the EXT1 L490Rfs*9 example,

endPositionInReferenceSequence = 490 + 9 - 2 = 497

The altered amino acid sequence itself is specified using the alteredAminoAcidFragment attribute. The end coordinate of the EWAS is changed to reflect the new C-terminal amino acid (Supplementary Methods Figure 5A).

If the research article does not completely describe all the details needed to annotate a frameshift variant, and if such details cannot be obtained from a disease variant database, they have to be deciphered.

The basic steps are:

1. Alter the reference coding sequence for the wild-type protein to match the coding sequence of the variant. The reference coding sequence can be obtained from the RefSeq [4] mRNA cross-referenced by the UniProt record that is used as the reference for the wild-type protein.
2. Translate the altered coding sequence in silico using EMBOSS TranSeq open software [5].
3. Align the resultant mutant peptide sequence with wild-type protein sequence using EMBOSS Needle [5] open software to determine the alteredAminoAcidFragment (and the name of the frameshift protein variant if not provided in the research article).

The disorder "Hereditary multiple exostoses 1" (EXT1) is caused by loss of function mutations in Exostosin 1 (EXT1). One mutation is a 1-bp deletion at the coding nucleotide 1469 in the EXT1 gene, resulting in a frameshift mutation with a premature stop codon [6,7]. The research articles, however, do not provide a name and details of the frameshift variant at the protein level. The reference UniProt record for the wild-type EXT1 protein in Reactome is Q16394, and it cross-references RefSeq mRNA NM_000127. The coding sequence in the RefSeq record starts at position 774. The nucleotide change, from literature, is a 1-bp thymidine deletion at the coding position 1469, which matches position 2242 (2242 = 774 - 1 + 1469) in NM_000127. The sequence surrounding the mutation should always be inspected to make sure that the coordinates provided in the paper are not outdated relative to RefSeq. Copy-paste the coding part of NM_000127, starting with the start codon, into EMBOSS Transeq Input Form, deleting the t at position 2242, immediately after the string of 6 cytidines and submit the job (Supplementary Methods Figure 5B).

The pairwise sequence alignment of the wild-type protein sequence and the EMBOSS Transeq output up to the first stop codon using EMBOSS Needle indicates the positions of the first altered amino acid and the premature stop codon, as well as the sequence of the altered fragment (Supplementary Methods Figure 5C).

**Frameshifts that produce a stop in the first affected codon** are annotated as nonsense mutations, as described above.

**Frameshifts that produce a stop codon in the second affected codon** are annotated as usual for a frameshift, as described above, with alteredAminoAcidFragment represented by a single letter.

##### Mutations affecting the initiator methionine

A mutation that eliminates the initiator methionine and results in the aberrant initiation of translation from a downstream methionine is annotated as an in-frame deletion, as per HGVS recommendations, with the variant protein name specifying a deletion that stretches from the second amino acid in the reference sequence to the alternative initiator methionine.

**Example:** SERPING1 A2_M53del - in this variant, a missense mutation substitutes the initiator methionine (M1) codon with a leucine codon, leading to the usage of methionine at position 53 (M53) as an alternative start codon.

**How to annotate:** FragmentDeletionModification specifies deletion of amino acid residues 1 to 52, and the protein variant EWAS record specifies the start coordinate as 53, without changing the end coordinate relative to the reference wild-type sequence (Supplementary Methods Figure 5C).

**If a gene translocation fuses the promoter of one gene with the coding sequence of another gene, with accompanying truncation of the N' terminal coding sequence of the C=terminal fusion partner,** the resulting disease variant is annotated as described here for mutations affecting the initiator methionine. The only such example in Reactome is the NOTCH1 variant NOTCH1 P2_M1580del (commonly known as TAN-1), which results from a translocation t(7;9) that fuses the promoter of T cell receptor beta to intron 24 of NOTCH1, resulting in the usage of an alternative start codon at position 1580 of the reference NOTCH1 sequence.

**Extension variants**

The only currently annotated extension variant is IKBKG *420Wext*447, which represents a 3' extension, with replacement of the stop codon at position 420 with tryptophan codon and extension of the protein sequence by a total of 27 amino acids, with a new stop codon at position 447, counting from the start codon of the reference wild-type protein sequence. Both 3' and 5' extensions can be annotated using fragmentReplacementModification class. In the case of 3' extensions, the wild-type stop codon coordinate is specified as the start and end coordinate of the replacement, and the added stretch of amino acids, in single letter abbreviations, is defined in the alteredAminoAcidFragment attribute (Supplementary Methods Figure 5D). In the case of 5' extensions, the -1 would be specified as the start and end coordinate of the replacement.

**Splicing variants,** if the exact effect of the mutation has been determined experimentally at the protein level, are annotated in one of the previously described categories, depending on the consequence of the splicing mutation on the amino acid sequence, and named accordingly, as HGVS currently does not provide guidelines for nomenclature of splicing variants at the protein level. This also applies to **inversion variants**. Of the 6 splicing variants currently annotated in Reactome, four are annotated as frameshift variants (EXT2 A361Pfs*44, NF1 Q616Gfs*4, NF1 R69Nfs*7, and UNC93B1 L230Afs*188), one is annotated as an in-frame insertion (POMT2 I444_N445insLLWQ), and one as an in-frame deletion (SLC11A2 L104_P142del). Two inversion variants currently annotated in Reactome, F8 A20_M2143delinsMRIQDPGK and F8 V2144_Y2351delinsHGVLENGKNWEPSYRW, are annotated as in-frame indels, as their names imply.

For **variants with multiple in-cis mutations**, which usually contain a primary mutation and a secondary mutation that was selected for during the course of treatment with target therapy as a drug-resistance mechanism, each mutation is annotated independently and they are separated with a semicolon in the disease variant name, e.g. EGFR E746_A750del;T790M, where T790M is a mutation associated with resistance to small tyrosine kinase inhibitors.

#### Cypher Query to Retrieve Disease Variant Information from Reactome GraphDB

```
MATCH ewas_to_reaction_path=(reaction_like_event:ReactionLikeEvent)-
[:input|output|catalystActivity|physicalEntity|regulatedBy|regulator|hasComponent|hasMem
ber|hasCandidate*]->(ewas:EntityWithAccessionedSequence)

WHERE ewas.speciesName="Homo sapiens"

WITH ewas,

    reaction_like_event,

    relationships(ewas_to_reaction_path) AS ewas_to_reaction_path_rels,

    nodes(ewas_to_reaction_path) AS ewas_to_reaction_path_nodes

MATCH

 (ewas:EntityWithAccessionedSequence)-[:referenceEntity]-
>(reference_gene_product:ReferenceGeneProduct),

   (ewas:EntityWithAccessionedSequence)-[:disease]->(disease:Disease),

   (ewas:EntityWithAccessionedSequence)-[:hasModifiedResidue]-
>(genetically_modified_residue:GeneticallyModifiedResidue),

   (reaction_like_event)-[:literatureReference]-
>(reaction_literature_reference:LiteratureReference),

   (reaction_like_event)<-[:hasEvent]-(disease_path:Pathway)

OPTIONAL MATCH (ewas:EntityWithAccessionedSequence)-[:referenceEntity]-
>(reference_entity:ReferenceEntity)-[:referenceDatabase]-
>(reference_entity_reference_database:ReferenceDatabase)//,

OPTIONAL MATCH (reaction_like_event)-[:normalReaction]-
>(normal_reaction_like_event:ReactionLikeEvent)

OPTIONAL MATCH (normal_reaction_like_event)-[:goBiologicalProcess]-
>(normal_reaction_like_event_go_biological_process:GO_BiologicalProcess)

OPTIONAL MATCH (disease_path)-[:normalPathway]->(pathway:Pathway)

OPTIONAL MATCH (pathway)-[:goBiologicalProcess]->(pgobp:GO_BiologicalProcess)
```

```
OPTIONAL MATCH (ewas:EntityWithAccessionedSequence)-[:literatureReference]->(literature_reference:LiteratureReference)

OPTIONAL MATCH (ewas:EntityWithAccessionedSequence)-[:crossReference]->(cross_reference:DatabaseIdentifier)

OPTIONAL MATCH (reaction_like_event)-[:entityFunctionalStatus]->(:EntityFunctionalStatus)-[:functionalStatus]->(:FunctionalStatus)-[:functionalStatusType]->(fst:FunctionalStatusType)

WITH ewas,
    reference_gene_product,
    reference_entity,
    reference_entity_reference_database,
    genetically_modified_residue,
    cross_reference,
    reaction_like_event,
    normal_reaction_like_event,
    disease.displayName AS disease_name,
    disease.databaseName + ":" + disease.identifier  AS disease_id,
    disease_path,
    pathway,
    normal_reaction_like_event_go_biological_process,
    pgobp,
    reaction_literature_reference,
    literature_reference,
    ewas_to_reaction_path_nodes,
    last([i IN range(0,size(ewas_to_reaction_path_nodes)-1) WHERE "CandidateSet" IN labels(ewas_to_reaction_path_nodes[i]) OR "EntitySet" IN labels(ewas_to_reaction_path_nodes[i]) | {node: ewas_to_reaction_path_nodes[i], relationship: type(ewas_to_reaction_path_rels[i])}]) AS first_entity_or_candidate_set_obj,
    fst
WITH ewas,
    reference_gene_product,
    reference_entity,
```

```
    reference_entity_reference_database,
    genetically_modified_residue,
    cross_reference,
    reaction_like_event,
    normal_reaction_like_event,
    disease_name,
    disease_id,
    disease_path,
    pathway,
    normal_reaction_like_event_go_biological_process,
    pgobp,
    reaction_literature_reference,
    literature_reference,
    ewas_to_reaction_path_nodes,
    CASE WHEN first_entity_or_candidate_set_obj.node.stId IS NULL THEN "" ELSE
    apoc.text.join([toString(first_entity_or_candidate_set_obj.node.stId),
                first_entity_or_candidate_set_obj.node.displayName,
                last(labels(first_entity_or_candidate_set_obj.node)),
                first_entity_or_candidate_set_obj.relationship], "-") END AS
first_distinct_entity_or_candidate_set,
    fst
WITH reference_gene_product.geneName[0] AS Genename,
   ewas.displayName AS displayName,
   ewas.stId AS stable_id,
   apoc.text.join(reference_entity.name,"|") AS referenceEntity_name,
   reference_entity.identifier AS referenceEntity_identifier,
   reference_entity_reference_database.displayName AS
referenceEntity_referenceDatabase_displayName,
   COLLECT(DISTINCT genetically_modified_residue.displayName) AS
hasModifiedResidue_displayName_array,
```

```
   COLLECT(DISTINCT genetically_modified_residue.schemaClass) AS
modifiedResidue_class_array,
   COLLECT(DISTINCT cross_reference.displayName) AS cross_reference_array,
   COLLECT(DISTINCT disease_name) AS disease_array,
   COLLECT(DISTINCT disease_id) AS disease_identifier_array,
   COLLECT(DISTINCT "pubmed:" + toString(literature_reference.pubMedIdentifier)) AS
entityWithAccessionedSequence_literatureReference_pubMedIdentifier_array,
   COLLECT(DISTINCT first_distinct_entity_or_candidate_set) AS first_entitySet,
   COLLECT(DISTINCT toString(reaction_like_event.stId)) AS
entityWithAccessionedSequence_reactionLikeEvent_stable_id_array,
   COLLECT(DISTINCT reaction_like_event.displayName) AS
entityWithAccessionedSequence_reactionLikeEvent_displayName_array,
   COLLECT(DISTINCT fst.displayName) AS
entityWithAccessionedSequence_reactionLikeEvent_entityFunctionalStatus_functionalStatus_fu
nctionalStatusType_displayName_array,
   COLLECT(DISTINCT "pubmed:" + toString(reaction_literature_reference.pubMedIdentifier)) AS
reactionLikeEvent_literatureReference_pubMedIdentifier_array,
   COLLECT(DISTINCT disease_path.stId) AS
entityWithAccessionedSequence_pathway_stable_id_array,
   COLLECT(DISTINCT disease_path.displayName) AS
entityWithAccessionedSequence_pathway_displayName_array,
   COLLECT(DISTINCT toString(normal_reaction_like_event.stId)) AS
entityWithAccessionedSequence_reactionLikeEvent_normalReaction_stable_id_array,
   COLLECT(DISTINCT normal_reaction_like_event.displayName) AS
entityWithAccessionedSequence_reactionLikeEvent_normalReaction_displayName_array,
   COLLECT(DISTINCT pathway.displayName) AS
entityWithAccessionedSequence_pathway_normalPathway_displayName_array,
   COLLECT(DISTINCT pathway.stId) AS
entityWithAccessionedSequence_pathway_normalPathway_stable_id_array,
   COLLECT(DISTINCT normal_reaction_like_event_go_biological_process.accession) AS
normal_reaction_like_event_go_biological_process_accession_array,
   COLLECT(DISTINCT normal_reaction_like_event_go_biological_process.displayName) AS
normal_reaction_like_event_go_biological_process_displayName_array,
```

```
   COLLECT(DISTINCT pgobp.accession) AS
entityWithAccessionedSequence_pathway_normalPathway_goBiologicalProcess_accession_arr
ay,

   COLLECT(DISTINCT pgobp.displayName) AS
entityWithAccessionedSequence_pathway_normalPathway_goBiologicalProcess_displayName_
array

RETURN DISTINCT

   Genename,

   displayName,

   stable_id,

   referenceEntity_name,

   referenceEntity_referenceDatabase_displayName + ":" + referenceEntity_identifier AS
referenceEntity_id,

   CASE WHEN size(hasModifiedResidue_displayName_array) = 0 THEN "" ELSE
apoc.text.join(hasModifiedResidue_displayName_array, "|") END AS
hasModifiedResidue_displayName,

   CASE WHEN size(modifiedResidue_class_array) = 0 THEN "" ELSE
apoc.text.join(modifiedResidue_class_array, "|") END AS modifiedResidue_class,

   CASE WHEN size(cross_reference_array) = 0 THEN "" ELSE
apoc.text.join(cross_reference_array, "|") END AS cross_reference,

   CASE WHEN size(disease_array) = 0 THEN "" ELSE apoc.text.join(disease_array, "|") END AS
disease,

   CASE WHEN size(disease_identifier_array) = 0  THEN "" ELSE
apoc.text.join(disease_identifier_array, "|") END AS disease_identifier,

   CASE WHEN
size(entityWithAccessionedSequence_literatureReference_pubMedIdentifier_array) = 0 THEN ""
ELSE
apoc.text.join(entityWithAccessionedSequence_literatureReference_pubMedIdentifier_array,
"|") END AS entityWithAccessionedSequence_literatureReference_pubMedIdentifier,

   CASE WHEN size(first_entitySet) = 0 THEN "" ELSE apoc.text.join(first_entitySet, "|") END AS
first_entitySet,

   CASE WHEN size(entityWithAccessionedSequence_reactionLikeEvent_stable_id_array) = 0
THEN "" ELSE
apoc.text.join(entityWithAccessionedSequence_reactionLikeEvent_stable_id_array, "|") END AS
entityWithAccessionedSequence_reactionLikeEvent_stable_id,
```

```
   CASE WHEN size(entityWithAccessionedSequence_reactionLikeEvent_displayName_array) = 0
THEN "" ELSE
apoc.text.join(entityWithAccessionedSequence_reactionLikeEvent_displayName_array, "|")
END AS entityWithAccessionedSequence_reactionLikeEvent_displayName,

   CASE WHEN
size(entityWithAccessionedSequence_reactionLikeEvent_entityFunctionalStatus_functionalStat
us_functionalStatusType_displayName_array) = 0 THEN "" ELSE
apoc.text.join(entityWithAccessionedSequence_reactionLikeEvent_entityFunctionalStatus_func
tionalStatus_functionalStatusType_displayName_array, "|") END  AS
entityWithAccessionedSequence_reactionLikeEvent_entityFunctionalStatus_functionalStatus_fu
nctionalStatusType_displayName,

   CASE WHEN size(reactionLikeEvent_literatureReference_pubMedIdentifier_array) = 0 THEN ""
ELSE apoc.text.join(reactionLikeEvent_literatureReference_pubMedIdentifier_array, "|") END
AS reactionLikeEvent_literatureReference_pubMedIdentifier,

   CASE WHEN size(entityWithAccessionedSequence_pathway_stable_id_array) = 0 THEN ""
ELSE apoc.text.join(entityWithAccessionedSequence_pathway_stable_id_array, "|") END AS
entityWithAccessionedSequence_pathway_stable_id,

   CASE WHEN size(entityWithAccessionedSequence_pathway_displayName_array) = 0 THEN ""
ELSE apoc.text.join(entityWithAccessionedSequence_pathway_displayName_array, "|") END AS
entityWithAccessionedSequence_pathway_displayName,

   CASE WHEN
size(entityWithAccessionedSequence_reactionLikeEvent_normalReaction_stable_id_array) = 0
THEN "" ELSE
apoc.text.join(entityWithAccessionedSequence_reactionLikeEvent_normalReaction_stable_id_a
rray, "|") END AS
entityWithAccessionedSequence_reactionLikeEvent_normalReaction_stable_id,

   CASE WHEN
size(entityWithAccessionedSequence_reactionLikeEvent_normalReaction_displayName_array)
= 0 THEN "" ELSE
apoc.text.join(entityWithAccessionedSequence_reactionLikeEvent_normalReaction_displayNam
e_array, "|") END AS
entityWithAccessionedSequence_reactionLikeEvent_normalReaction_displayName,

   CASE WHEN
size(entityWithAccessionedSequence_pathway_normalPathway_displayName_array) = 0 THEN
"" ELSE
apoc.text.join(entityWithAccessionedSequence_pathway_normalPathway_displayName_array,
"|") END AS entityWithAccessionedSequence_pathway_normalPathway_displayName,

   CASE WHEN
size(entityWithAccessionedSequence_pathway_normalPathway_stable_id_array) = 0 THEN ""
ELSE
```

```
apoc.text.join(entityWithAccessionedSequence_pathway_normalPathway_stable_id_array, "|")
END AS entityWithAccessionedSequence_pathway_normalPathway_stable_id,

   CASE WHEN size(normal_reaction_like_event_go_biological_process_accession_array) = 0
THEN "" ELSE
apoc.text.join(normal_reaction_like_event_go_biological_process_accession_array, "|") END AS
normal_reaction_like_event_go_biological_process_accession,

   CASE WHEN size(normal_reaction_like_event_go_biological_process_displayName_array) = 0
THEN "" ELSE
apoc.text.join(normal_reaction_like_event_go_biological_process_displayName_array, "|")
END AS normal_reaction_like_event_go_biological_process_displayName,

   CASE WHEN
size(entityWithAccessionedSequence_pathway_normalPathway_goBiologicalProcess_accession
_array) = 0 THEN "" ELSE
apoc.text.join(entityWithAccessionedSequence_pathway_normalPathway_goBiologicalProcess_
accession_array, "|") END AS
entityWithAccessionedSequence_pathway_normalPathway_goBiologicalProcess_accession,

   CASE WHEN
size(entityWithAccessionedSequence_pathway_normalPathway_goBiologicalProcess_displayNa
me_array) = 0 THEN "" ELSE
apoc.text.join(entityWithAccessionedSequence_pathway_normalPathway_goBiologicalProcess_
displayName_array, "|") END AS
entityWithAccessionedSequence_pathway_normalPathway_goBiologicalProcess_displayName

ORDER BY Genename ASC
```

### Supplementary Methods Figure Legends

#### Supplementary Methods Figure 1.

Flowchart of **A)** manual and **B)** automated annotation of disease variants in Reactome.

#### Supplementary Methods Figure 2.

**A)** Overview of disease-related attributes annotated in disease variant records. **B)** Overview of GeneticallyModifiedResidue class used to describe genetically encoded amino acid changes in disease variants. **C)** Annotation of missense mutations.

#### Supplementary Methods Figure 3.

**A)** Annotation of nonsense variants. **B)** Annotation of in-frame deletion variants that lack multiple consecutive amino acids. **C)** Annotation of in-frame deletion variants that lack a single amino acid. **D)** Annotation of in-frame insertion variants.

#### Supplementary Methods Figure 4.

Annotation of **A)** duplication variants; **B)** fusion variants; **C)** fusion variants with translated in-frame intronic/UTR insertions; **D)** in-frame indel variants.

#### Supplementary Methods Figure 5.

**A)** Annotation of frameshift protein variants. **B)** Inference of amino acid sequence change in protein frameshift variants from change in the coding sequence of the reference transcript. **C)** Annotation of mutations in the initiator methionine codon that lead to the usage of an alternative translation start. **D)** Annotation of 3’ extension variants.

### Supplementary Methods Figure 1.

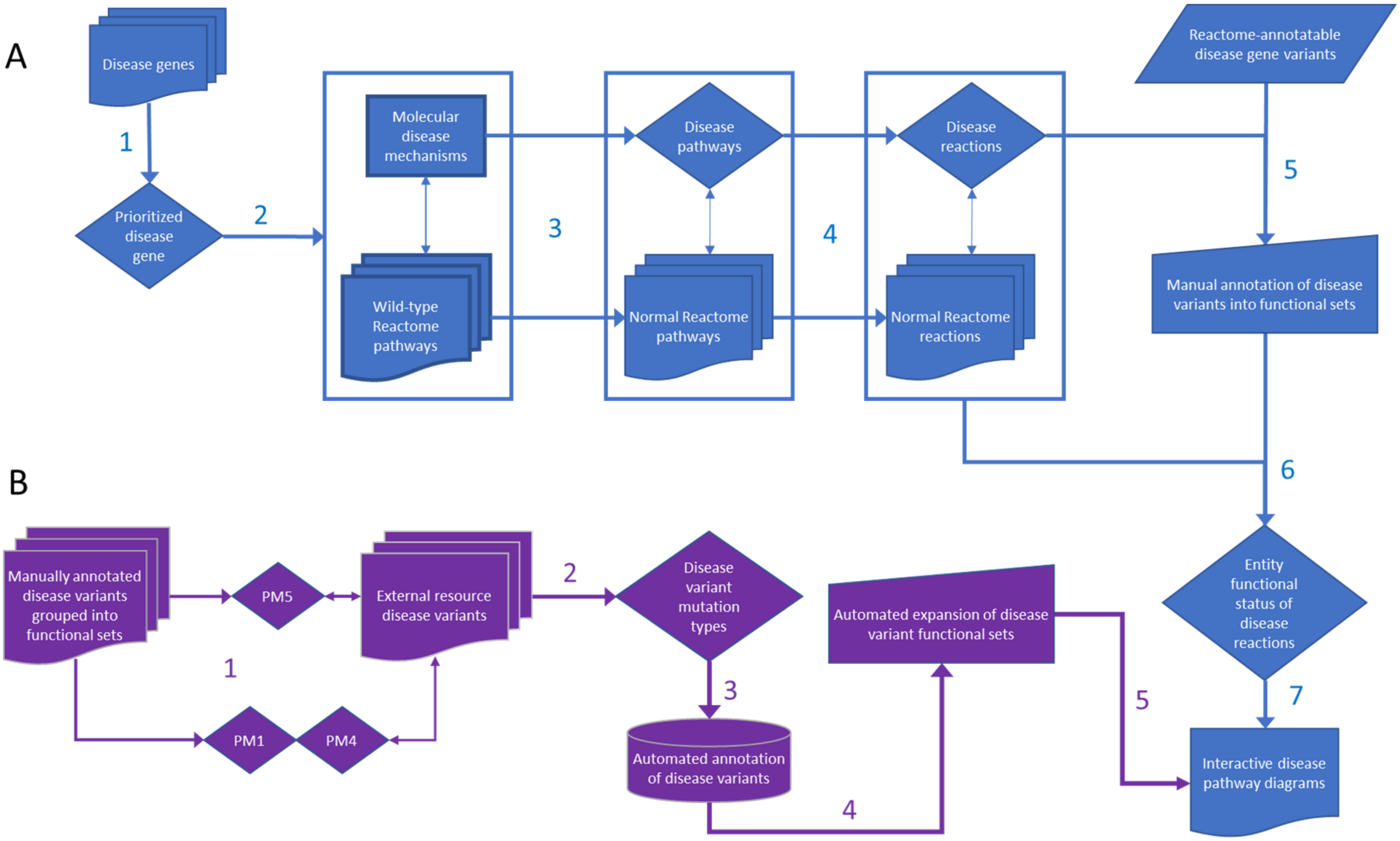

### Supplementary Methods Figure 2.

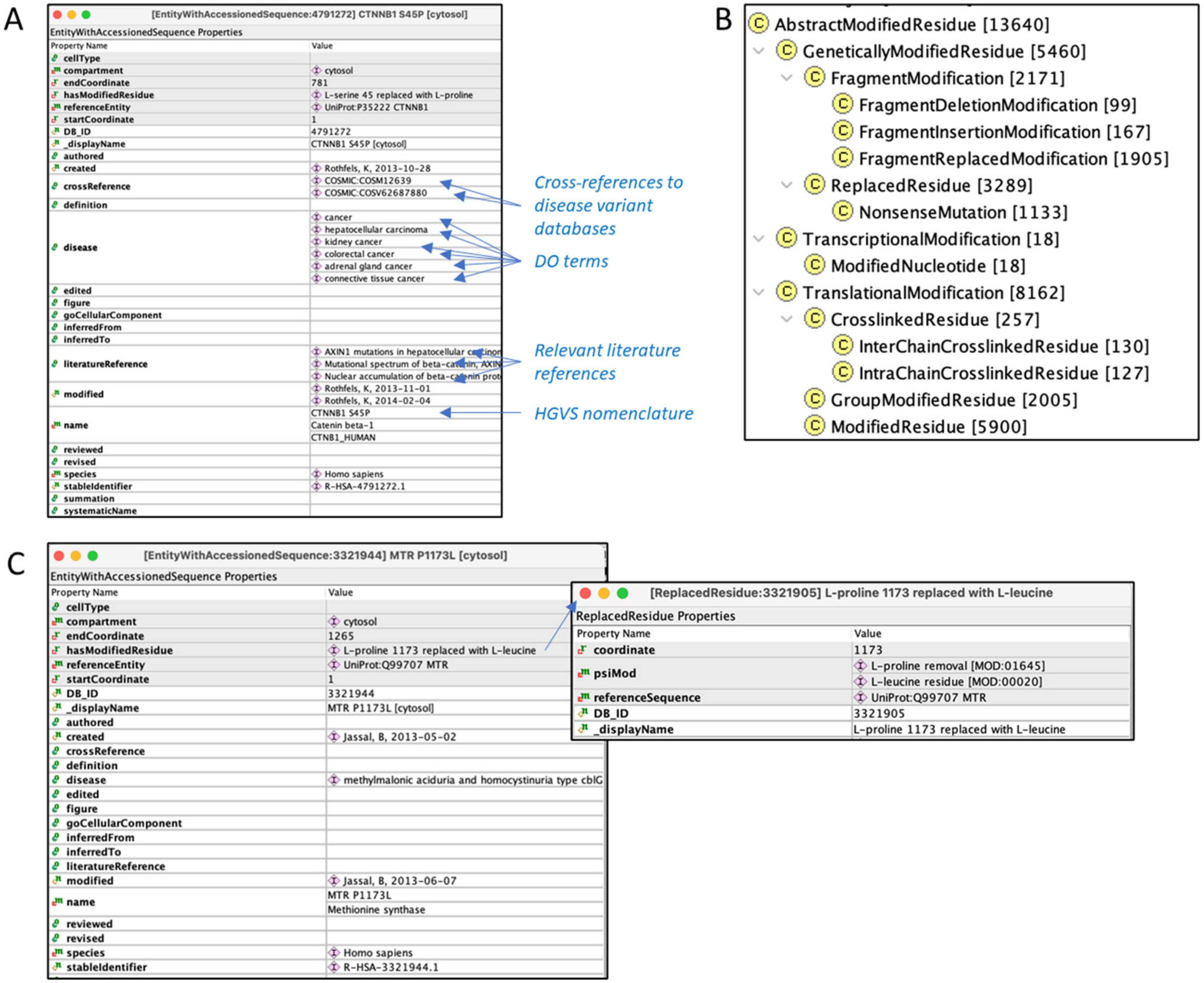

### Supplementary Methods Figure 3.

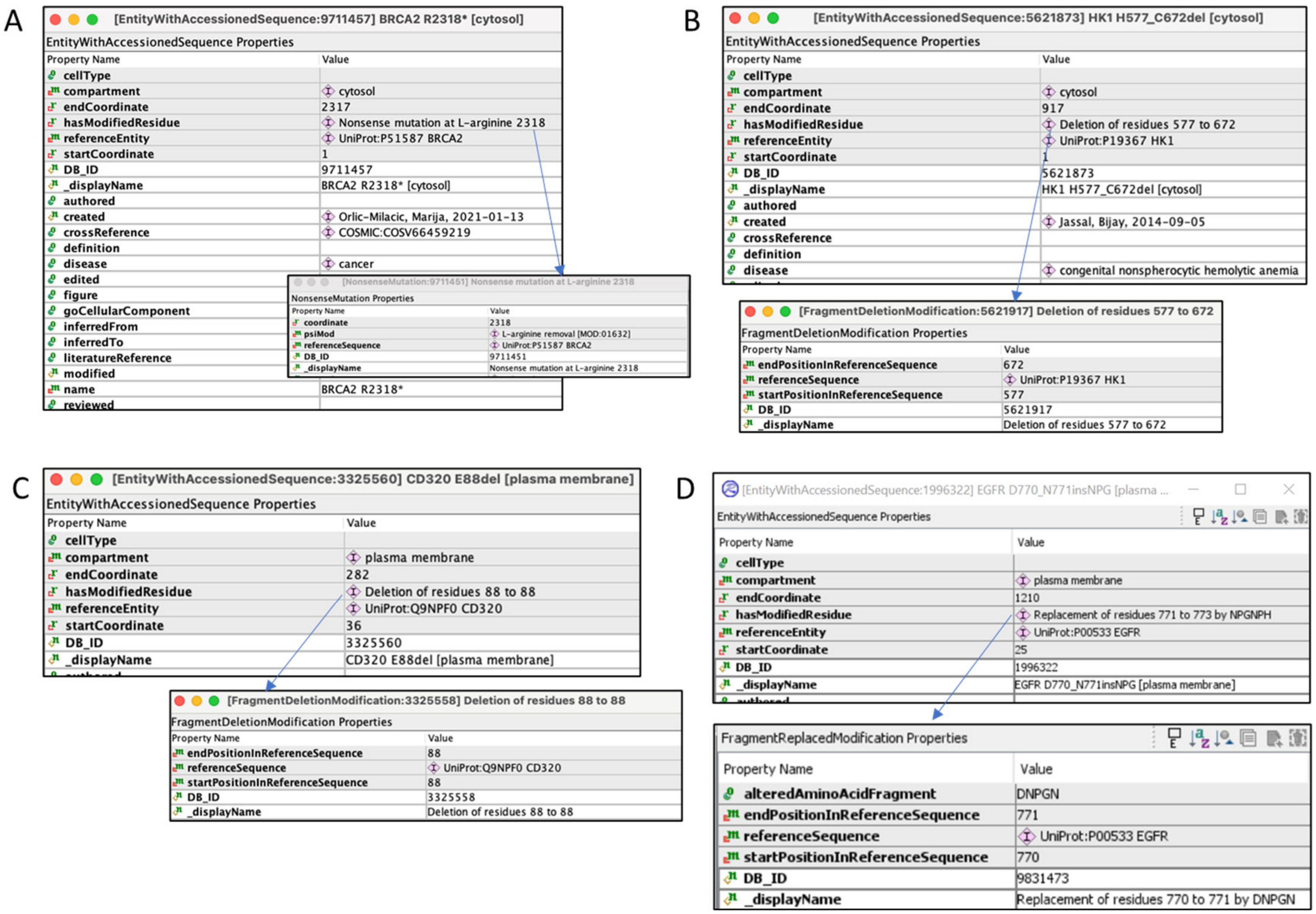

### Supplementary Methods Figure 4.

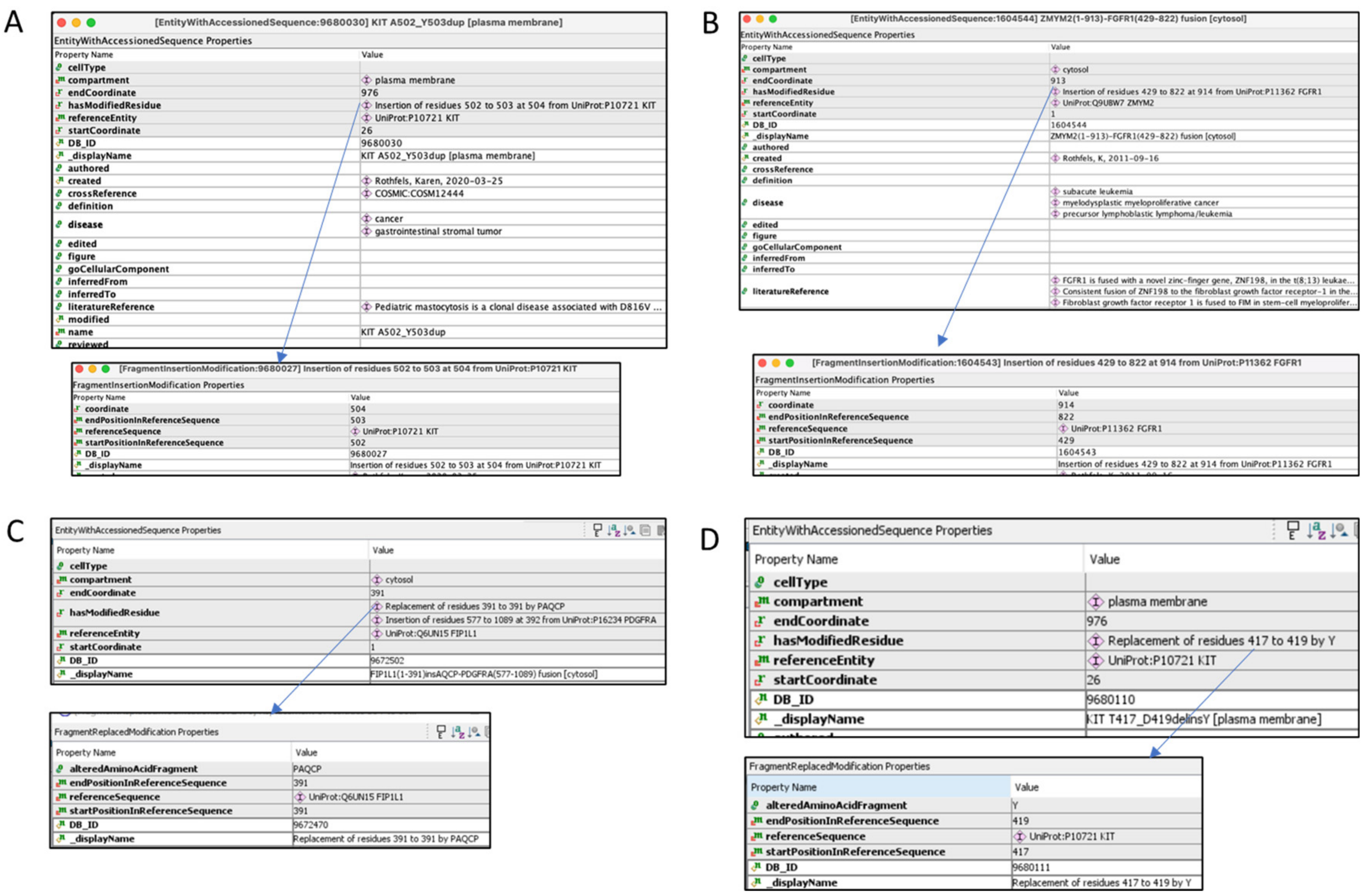

### Supplementary Methods Figure 5.

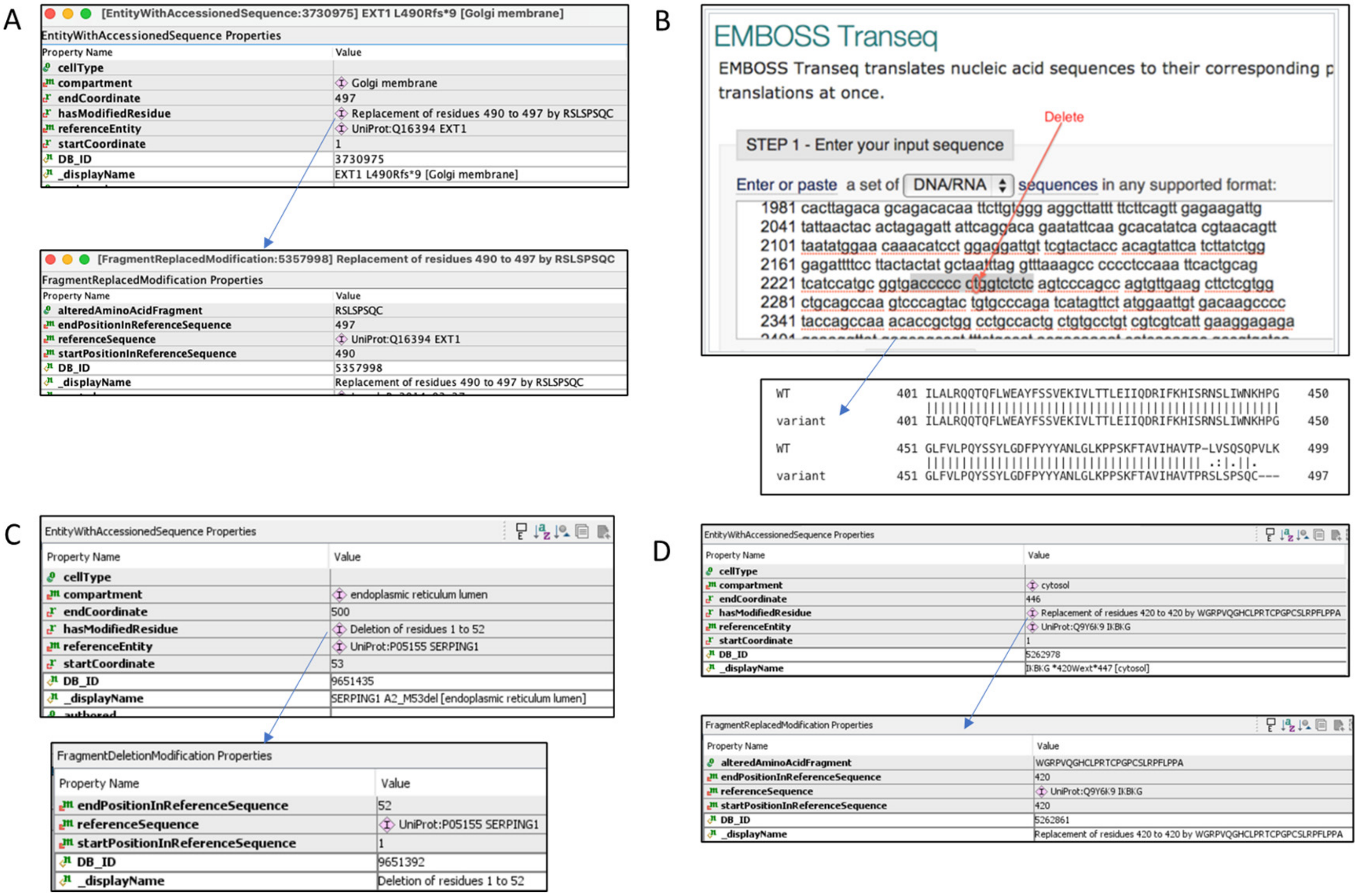
